# Structural insights into Measles virus RNA synthesis: C protein–mediated regulation of replication-transcription switch and allosteric inhibition by ERDRP-0519

**DOI:** 10.1101/2025.07.21.666041

**Authors:** Tianjiao Du, Jiening Wang, Chengji Yang, Guobao Li, Kaiyue Jie, Ying Chen, Xiaokang Zhang, Long Zhang, Gaojie Song, Qiansen Zhang, Shan Wu, Heng Ru

**Author notes:** These authors contributed equally: Tianjiao Du, Jiening Wang and Chengji Yang. These authors jointly supervised this work: Qiansen Zhang, Shan Wu, Heng Ru.

## Abstract

Nonsegmented negative-sense RNA viruses (nsNSVs) rely on a multifunctional RNA-dependent RNA polymerase (RdRP) complex for transcription and replication. In measles virus (MeV), the nonstructural protein C has long been implicated in regulating RNA synthesis, yet its precise role remains unclear. Here, we show that the MeV C protein directly associates with the RdRP complex. Using cryo-electron microscopy, we determined atomic-resolution structures of the MeV polymerase with and without C, revealing that C binding stabilizes the accessory domains of L and locks the complex into a replication-competent elongation state. Biochemical data further show that C promotes N protein recruitment, enhancing polymerase processivity through facilitating encapsidation during replication. Additionally, we also resolved high-resolution structures of MeV and Nipah virus (NiV) polymerases bound to ERDRP-0519, an orally available morbillivirus inhibitor. Unexpectedly, the compound occupies a previously unrecognized allosteric pocket within the RdRp domain, rather than the predicted PRNTase domain, overlapping conserved resistance sites. This binding induces conformational changes in palm subdomain, blocking RNA template and nucleotide engagement, thereby halting RNA synthesis. These findings uncover distinct regulatory and inhibitory mechanisms in paramyxovirus polymerases and provide a structural framework for the rational design of broad-spectrum antivirals targeting MeV, NiV, and potentially other clinically relevant nsNSVs.

## INTRODUCTION

Nonsegmented, negative-strand RNA viruses (nsNSVs) constitute a large and diverse population of pathogenic viruses, including members of the *Paramyxoviridae*, *Pneumoviridae*, *Filoviridae*, *Rhabdoviridae*, *Bornaviridae* families, etc^1, 2^. These viruses cause a broad spectrum of diseases in humans and other mammals, ranging from mild infections to severe and often fatal illnesses, thereby posing a significant threats to public health and the livestock industry^1, 2^. The genomes of nsNSVs consist of linear, single-stranded and negative-sense RNA, encapsidated by nucleocapsid proteins (NPs) to form filamentous ribonucleoprotein (RNP) complexes, which serve as templates for both transcription and replication^1, 2, 3, 4^. The genomes are typically 11 to 19 kb in length and are flanked by a 3′ leader and a 5′ trailer sequence, both functioning as promoters for RNA synthesis^1, 2, 3, 4^. The viral genes are arranged sequentially and transcribed in tandem, each flanked by *cis*-acting gene start (GS) and gene end (GE) signals within their untranslated regions (UTRs), which are essential for regulating viral RNA synthesis^1, 2, 3, 4^. Upon entry into the host cell and release of viral RNPs into the cytoplasm, the nsNSV polymerase initiates transcription at the 3′ promoter. Following synthesis and release of a short non-coding RNA corresponding to the promoter-proximal leader region, the polymerase remains engaged with the template to scan for the GS signal of the first gene. Transcription of each gene produces a capped and methylated monocistronic mRNA, which is elongated until a GE signal is encountered, triggering polyadenylation and release of the transcript. The polymerase then resumes scanning and reinitiates transcription at the next downstream GS signal. This transcriptional strategy, known as the ‘stop–start’ mechanism, leads to a positional gradient of mRNA abundance, with genes proximal to the 3′ leader (e.g., NP) transcribed more frequently, whereas genes near the 5′ trailer (e.g., L), are produced at significantly lower levels due to polymerase attenuation at gene junctions^2, 3, 4^. In contrast, replication also initiates at the 3′ end of the viral genome but the polymerase proceeds with high processivity along the entire template without responding to internal *cis*-acting signals. This results in the synthesis of a full-length, uncapped, and non-polyadenylated positive-sense antigenome, which serves as a faithful complementary template for generating progeny genomes. Continuous encapsidation of nascent viral RNA by NPs is essential for productive replication, promoting functional RNP assembly and shielding the viral RNAs from host innate immune recognition^2, 3, 4^.

Viral RNA synthesis in nsNSVs is mediated by the RNA-dependent RNA polymerase (RdRP) complex, which comprises the large (L) protein and a cofactor—phosphoprotein (P) in most nsNSVs or VP35 in filoviruses^1, 2, 3, 4^. The L protein harbors all enzymatic activities essential for transcription and replication, including ribonucleotidyl transferase activity, polyribonucleotidyl transferase (PRNTase) activity for mRNA capping, and S-adenosylmethionine (SAM)-dependent N^7^-guanosine- and 2′-O-methyltransferase (MTase) activity^4, 5^. The cofactor P or VP35 is thought to maintain NP in a monomeric, RNA-free state (N_0_P) and facilitate its delivery to the polymerase complex, thereby ensuring efficient encapsidation of nascent RNA and supporting polymerase processivity during replication^2, 6^. Although the nsNSV RdRP complex carries out both transcription and replication, the molecular mechanism underlying the switch between these two mechanistically distinct processes and their regulation remain poorly understood. Several viral-encoded nonstructural proteins and host factors have been reported to play a role in the regulation of these processes. For example, in vesicular stomatitis virus (VSV)-infected cells, two distinct pools of RNA polymerase complexes, the transcriptase and replicase, have been purified, both containing the L–P complex associated with different components, with translation elongation factor-1a and heat shock protein 60 specifically identified in the transcriptase^7^. In addition, the M2-1 protein of human respiratory syncytial virus (RSV) acts as a transcription processivity factor, enabling the polymerase to read through the GE signals and the intragenic regions to transcribe the full set of viral genes^8, 9, 10, 11^. Furthermore, Ebolavirus (EBOV) VP30 functions as a transcription activator that initiates viral mRNA synthesis by resolving an inhibitory RNA hairpin at the NP gene start site, with its activity tightly regulated by phosphorylation^12, 13, 14, 15, 16^, whereas Marburgvirus (MARV) VP30 which shares high structural similarity with EBOV VP30, influences both transcription and replication, and is essential for the viral life cycle, yet appears dispensable for transcription activation in cell-based minigenome assays^13, 16, 17, 18, 19^. In contrast, in several viruses, including Sendai virus (SeV), Measles virus (MeV), Nipah virus (NiV), and human parainfluenza virus type 3 (HPIV3), the L, P, and N genes are sufficient to support reporter gene expression in minigenome assays, suggesting that the transcription of viral mRNAs does not rely on additional viral factors^20, 21, 22, 23^. The C protein, a nonstructural product translated from an alternative initiation site on the P mRNA, is known to act as a virulence factor by antagonizing host innate immune signaling pathways^24^. During infection with MeV, SeV, and HPIV1, the absence of C protein expression strongly induces the production of non-encapsidated copy-back defective interfering RNAs (cbDI-RNAs), which potently activate the innate immune response and can consequently restrict viral propagation^25, 26, 27, 28, 29, 30, 31, 32, 33^. In addition to its immunomodulatory role, the C protein is also thought to enhance the processivity of the viral RNA polymerase during replication; however, whether it acts directly on the polymerase and the precise mechanism underlying this regulation remain unclear.

As representative members of the morbillivirus and henipavirus genera within the *Paramyxoviridae* family, MeV and NiV are highly contagious respiratory pathogens that typically cause acute disease and pose a significant threats to human health^34, 35^. MeV is transmitted via respiratory route and initially presents with fever, cough, coryza, and conjunctivitis, followed by the appearance of a characteristic rash. Complications may affect multiple organ systems, with pneumonia and subacute sclerosing panencephalitis being the major contributors to measles-associated morbidity and mortality^36, 37^. Although measles is a vaccine-preventable disease, it continues to impose a substantial global disease burden, causing an estimated 100,000 deaths annually. This persistence is attributed to declining vaccination coverage below the 95 percent threshold required for herd immunity, enabling the reestablishment of endemic transmission and driving widespread outbreaks^36, 37, 38, 39, 40, 41, 42^. NiV, a recently emerged zoonotic paramyxovirus, is transmitted through contact with infected animals or via human-to-human transmission. In humans, NiV infection causes systemic vasculitis that leads to symptoms such as fever, headache, dizziness, and confusion, which can rapidly progress to fatal encephalitis, with case fatality rates ranging from 40 to 90 percent^35, 43, 44, 45, 46^. Due to its broad host range, high pathogenicity, and the absence of approved vaccines or antiviral therapies, NiV represents a particularly serious pandemic threat^47^, underscoring the urgent need for effective antiviral drug development against such high-risk pathogens.

Due to the absence of a cellular counterpart and the fundamental differences between host and viral RNA synthesis mechanisms, viral RdRP complexes are considered promising targets for the development of antiviral drugs. Previous cell-based high-throughput screening (HTS) and structure–activity relationship (SAR) studies led to the discovery of a series of non-nucleoside inhibitors with a shared core scaffold, including 16677, AS-136a, and ERDRP-0519, all of which exhibit potent antiviral activity against MeV, with subnanomolar EC_50_ values in cell-based assays, and directly inhibit the RNA synthesis activity of the MeV polymerase^48, 49, 50, 51, 52^. Importantly, they are effective against a broad spectrum of wild-type MeV genotypes that are currently circulating globally^48, 49, 51^. Among them, ERDRP-0519 stands out due to its favorable water solubility and high oral bioavailability, making it a particularly promising candidate for therapeutic development^52^. Additionally, ERDRP-0519 has demonstrated strong prophylactic and therapeutic efficacy in animal models, protecting ferrets from otherwise 100% lethal canine distemper virus (CDV) infection and preventing measles disease in squirrel monkeys, thereby confirming its broad activity against morbilliviruses and potential for clinical use^53, 54^. A combination of biochemical and computational studies has shown that ERDRP-0519 uniquely inhibits MeV polymerase activity by blocking all phosphodiester bond formation—including *de novo* initiation and primer-based extension—through simultaneously engaging the PRNTase domain and a flexible intrusion loop within the L protein, thereby locking the polymerase in a pre-initiation conformation^55, 56^. However, resistance mutations predominantly cluster in the RdRp domains of the L proteins in both MeV and CDV^48, 55^, raising unresolved questions about the precise binding mode and inhibitory mechanism of the compound. Thus, structural elucidation is required to clarify how ERDRP-0519 suppresses RNA synthesis and whether its inhibitory scope can extend to other paramyxoviruses or even broader nsNSVs. GHP-88309, another non-nucleoside compound identified through HTS, has been reported to inhibit the polymerase activity of multiple paramyxoviruses, including MeV, CDV and HPIV3, and to confer resistance mutations that map to a conserved cavity in the L protein distinct from the ERDRP-0519 resistance sites^54, 57^, raising the question of whether GHP-88309 exerts its antiviral effect through a different mechanism or modulation of polymerase function.

In this study, we employed biochemical approaches and cryo-electron microscopy (cryo-EM) to investigate the structure and regulation of the MeV RdRP complex. We determined high-resolution cryo-EM structures of the MeV RdRP complex in both its apo form and in complex with the viral nonstructural protein C (LPC). Both structures adopt an elongation conformation, However, comparative analysis reveals that C binding stabilizes the C-terminal accessory domains of the L protein. Moreover, the LPC structure captures the polymerase in a replication-competent elongation rather than a transcriptional state. Biochemical analysis further shows that the C protein preferentially associates with monomeric and single-stranded RNA-bound N protein, but not with the N^0^P complex, suggesting a previously unrecognized role for C in recruiting N during replication. These findings expand the known regulatory functions of C, implicating it not only in modulating the transcription-replication switch but also in enhancing the processivity of the polymerase during genome synthesis.

Additionally, we resolved the cryo-EM structure of the MeV RdRP complex bound to the small-molecule inhibitor ERDRP-0519 at 3.1 Å resolution. Contrary to prior assumptions, ERDRP-0519 does not bind the PRNTase domain but instead occupies a narrow pocket formed by the fingers and palm subdomains within the RdRp domain. This site overlaps with known resistance mutations and is highly conserved across the paramyxovirus family, supporting the potential of ERDRP-0519 as a broad-spectrum antiviral. Furthermore, we demonstrated that ERDRP-0519 binds the NiV polymerase with comparable competence and determined the structure of the NiV polymerase–inhibitor complex at near-atomic resolution. As with MeV, inhibitor binding induces major conformational rearrangements in the palm subdomain, particularly in the loop containing the conserved GDNE motif, which disrupts the active site architecture. The inhibitor also creates steric clashes with the template, product RNA, and incoming nucleotide in the elongation state, thereby revealing the molecular basis for its inhibition of RNA synthesis.

Together, our findings provide critical mechanistic insights into viral RNA synthesis and its regulation by both viral cofactors and small-molecule inhibitors, offering a structural framework for the rational design and optimization of antiviral therapeutics targeting MeV, NiV, and a broad spectrum of viruses within and beyond the paramyxovirus family.

## RESULTS

### Cryo-EM structure determination of RdRP complexes of human measles virus

Initial attempts to reconstitute the full-length MeV L and P protein yielded heterogeneous assemblies, as indicated by a broad peak in size-exclusion chromatography (SEC) profiles (Supplementary Fig. 1A–C). To improve complex stability and homogeneity, we truncated the predicted intrinsically disordered N-terminal region of the P protein preceding its tetramerization domain. This engineering resulted in a well-behaved, monodisperse RdRP complex, as confirmed by SEC (Supplementary Fig. 1A, B, and D). The truncated P protein construct was thus used for all subsequent structural and biochemical studies. Using this optimized construct, we reconstituted three distinct RdRP complexes: the apo L–P complex, and L–P complexes bound to the allosteric inhibitors ERDRP-0519 and GHP-88309, respectively.

To assemble the LPC complex, we co-expressed MeV L, P, and C proteins in Sf9 insect cells, using excess amounts of P and C relative to L. Affinity purification via a C-terminal Strep II tag on the L protein co-eluted the C protein together with the L–P complex (Supplementary Fig. 1E). The three proteins also co-migrated as a single peak on gel-filtration chromatography, with an elution volume comparable to that of the L–P complex (Supplementary Fig. 1B and F), indicating the formation of a stable trimeric complex under near-physiological conditions. To determine whether C associates with L or P, we performed a reverse pulldown using a C-terminal GFP-tagged C protein and anti-GFP nanobody to enrich binding partners from the flow-through fraction of the Strep affinity purification. Despite an excess of P protein in the input, only minimal amounts were recovered with C, suggesting limited C–P interaction (Supplementary Fig. 1G). Furthermore, purified C protein failed to bind either the full-length or N-terminally truncated P protein used in our structural studies (Supplementary Fig. 1H). These findings, in contrast to earlier reports^58^, strongly support a direct interaction between C and L proteins. Consistently, microscale thermophoresis (MST) analysis confirmed a high-affinity interaction between C and the L–P complex, with a dissociation constant (*K*_d_) of approximately 320 nM (Supplementary Fig. 1I).

All complexes, including the apo L–P complex, L–P complex bound to non-structure protein C (hereafter referred to as the LPC complex), and L–P complexes with inhibitors ERDRP-0519 and GHP-88309, were purified to homogeneity and subjected to cryo-EM sample preparation. Multiple datasets were collected for each complex on an FEI Titan Krios transmission electron microscope equipped with a K3 camera (Supplementary Table 1). Cryo-EM structure determination using multiple rounds of 2D, 3D classification and refinement yields a number of density maps, including the apo L–P complex, the LPC complex at 2.7 Å resolution, and the ERDRP-0519-bound complex at 3.1 Å resolution (Supplementary Fig. S2-4). Unexpectedly, we observed no discernible electron density corresponding to GHP-88309 in the L–P–GHP-88309 complex, despite increasing the compound concentration in a second dataset. The resulting map was nearly indistinguishable from that of the apo L–P complex (Supplementary Fig. S4B). Therefore, particles from the apo-form, the GHP-88309 datasets, and the LPC datasets lacking bound C protein were pooled and used to generate a composite apo-form map at 2.6 Å resolution (Supplementary Fig. S4). The map Fourier shell correlation (FSC) curves and local resolution estimations are in accordance with the gold-standard resolutions between half maps of split data.

In the L–P complex, the cryo-EM density map resolves the RdRp and PRNTase domains of the L protein, whereas the C-terminal regions, including the connector domain (CD), methyltransferase (MTase) domain, and C-terminal domain (CTD), are not visible (Fig. 1A and G), despite the L protein being intact during sample preparation (Supplementary Fig. 1E). This absence likely reflects intrinsic flexibility of the C-terminal region relative to the polymerase core under the current conformational state. The P protein forms a tetramer via its oligomerization domain, and all four subunits stably associate with the RdRp domain of L (Fig. 1A and G). In contrast, the LPC complex reveals nearly complete density for the L protein, including its full C-terminal region, except for several flexible loops (Fig. 1B, C and G). Notably, two C protein molecules, which associate with each other, are observed contacting both the RdRp/PRNTase core and the C-terminal domains of L, occupying a position above the polymerase central cavity. The ERDRP-0519-bound RdRP complex map closely resembles the apo L–P complex but features a prominent additional density within a deep cleft of the RdRp domain (Fig. 1D and E). This density is absent in both the L–P and LPC complexes and is attributed to ERDRP-0519 binding. These high-resolution reconstructions allowed unambiguous assignment of secondary structures and side-chain orientations. Atomic models were built and iteratively refined in real and reciprocal space, showing excellent agreement with their respective maps (Supplementary Fig. 5). Structural superposition of all three models revealed highly conserved configurations in the L protein core and P tetramers (Fig. 1F).

**Fig. 1.**
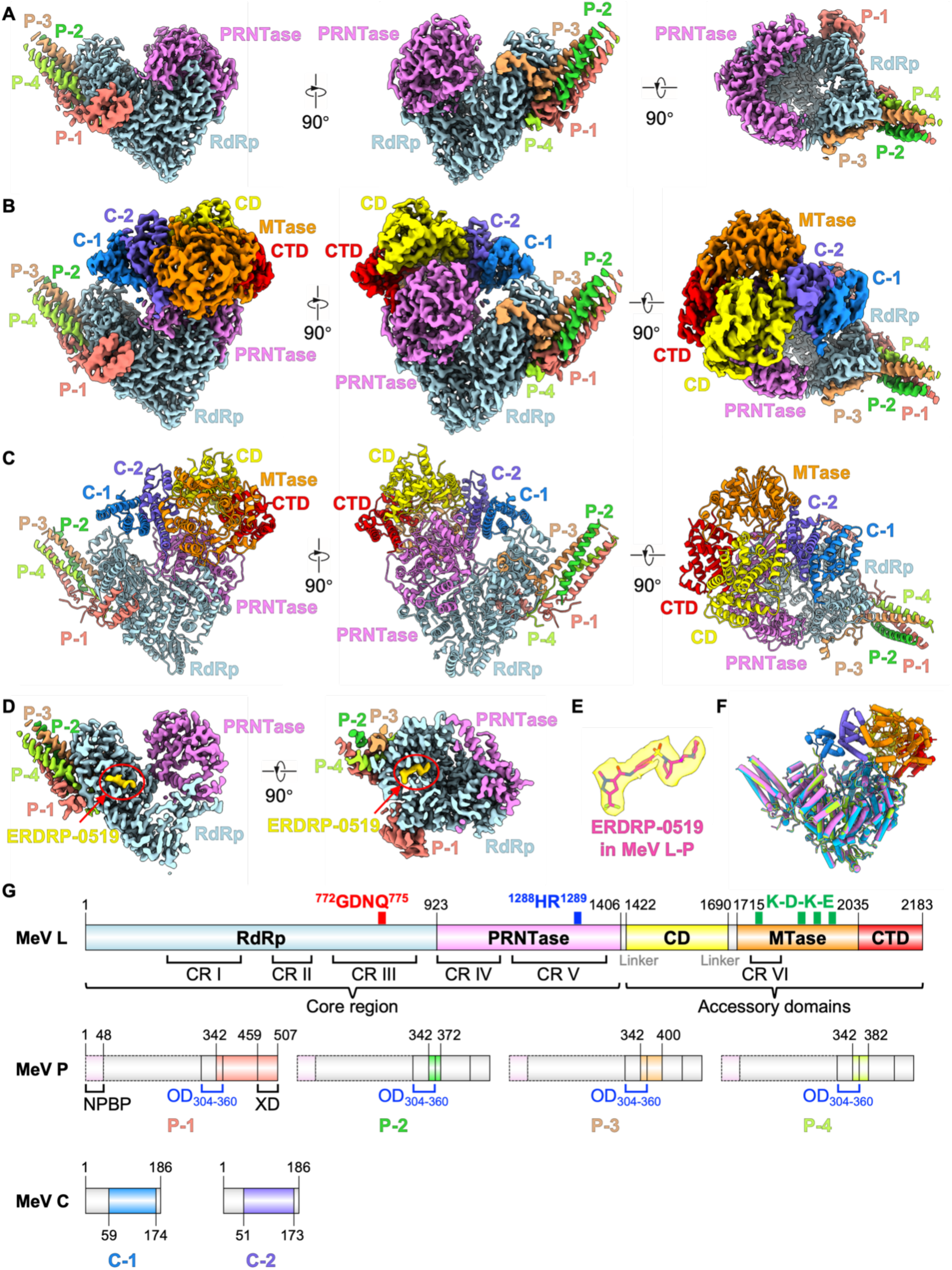
Overview of the cryo-EM maps and models of MeV RdRP complexes. **A**, Orthogonal views of the cryo-EM map of the MeV apo L–P complex. The resolved domains of the L protein, including the RdRp and PRNTase domains, are labeled and shown in light blue and pink, respectively. Four P protomers with varying lengths are visualized and colored as follows: P-1 in salmon, P-2 in lime green, P-3 in wheat, and P-4 in limon. **B and C**, Orthogonal views of the cryo-EM map and model of the MeV L–P complex bound to the C protein dimer (LPC complex). Upon C protein engagement, the accessory domains of the L protein become visible, including the connector domain (CD, yellow), methyltransferase domain (MTase, orange), and C-terminal domain (CTD, red). The two C protomers are shown in marine and slate, respectively. The core region of L and the P protomers are colored as in **A**. **D**, Cross-sectional view of the cryo-EM map of the L–P complex bound to ERDRP-0519 (LPE complex). The density corresponding to ERDRP-0519 is highlighted in yellow within the RdRp domain. **E**, Zoomed-in view of the ERDRP-0519 density in the cryo-EM map superimposed with the fitted atomic model. **F**, Superposition of the MeV RdRP complex models: apo L–P (pink), LPE (blue), and LPC (light green). The accessory domains and C protein dimers are shown in the same colors as in **B** and **C**. **G**, Schematic domain organization of the MeV L, P, and C proteins. Conserved regions I–VI (CR I–VI) in the L protein are labeled based on sequence alignment across paramyxoviruses. Catalytic residues within the RdRp, PRNTase, and MTase domains are annotated. Domain colors match those used in the cryo-EM maps and models. Gray boxes with solid outlines indicate unresolved regions in the structures, while gray boxes with dashed outlines represent regions that are not included in the constructs.

### Structural features of the RdRp and PRNTase domains in the MeV L protein

Among the three structures, the apo L–P complex exhibits the highest resolution and was thus used to describe the structural features of the L protein. The resolved core spans residues from 7 to 1405 and comprises two domains, the RdRp and PRNTase domain, which interlock to form a doughnut-shaped architecture with a central cavity (Fig. 1A), a conserved feature observed across nsNSVs. Structural comparison of the MeV L core with other *Paramyxoviridae* L proteins yields root mean square deviations (r.m.s.d.) of ∼1.2–2.1 Å, underscoring their evolutionary conservation within the family (Supplementary Fig. S6A–E), while indicating more distant relationships with L proteins from other *Mononegavirales* families. (Supplementary Fig. S6F–L). Analogous to the other polymerases, the RdRp domain adopts a canonical right-handed fingers–palm–thumb configuration (Fig. 2A), comprising seven conserved sequence motifs (motif A–G) (Fig. 2B). A globular N-terminal domain (NTD) precedes the fingers and likely serves as a structural scaffold. The highly conserved ^772^GDNQ^775^ motif, which is the characteristic of the catalytic residues in the RdRp domain of L protein among the nsNSVs, locates in the motif C of the palm subdomain. This motif, together with the conserved D663 residue in motif A, forms the active site of the RdRp domain (Fig. 2B), which was proposed to make phosphodiester bond via a “two-metal catalysis” mechanism. Although magnesium ion was present throughout the purification, no metal density was observed near the active site, suggesting that proper catalytic configuration may require substrate binding. AlphaFold3 predictions indicate a disordered insertion between the supporting helix and motif A (Supplementary Fig. 6M). In our structure, this region including the supporting helix itself is unresolved, likely due to its high flexibility in the apo conformation, preventing stable association with the palm subdomain.

**Fig. 2.**
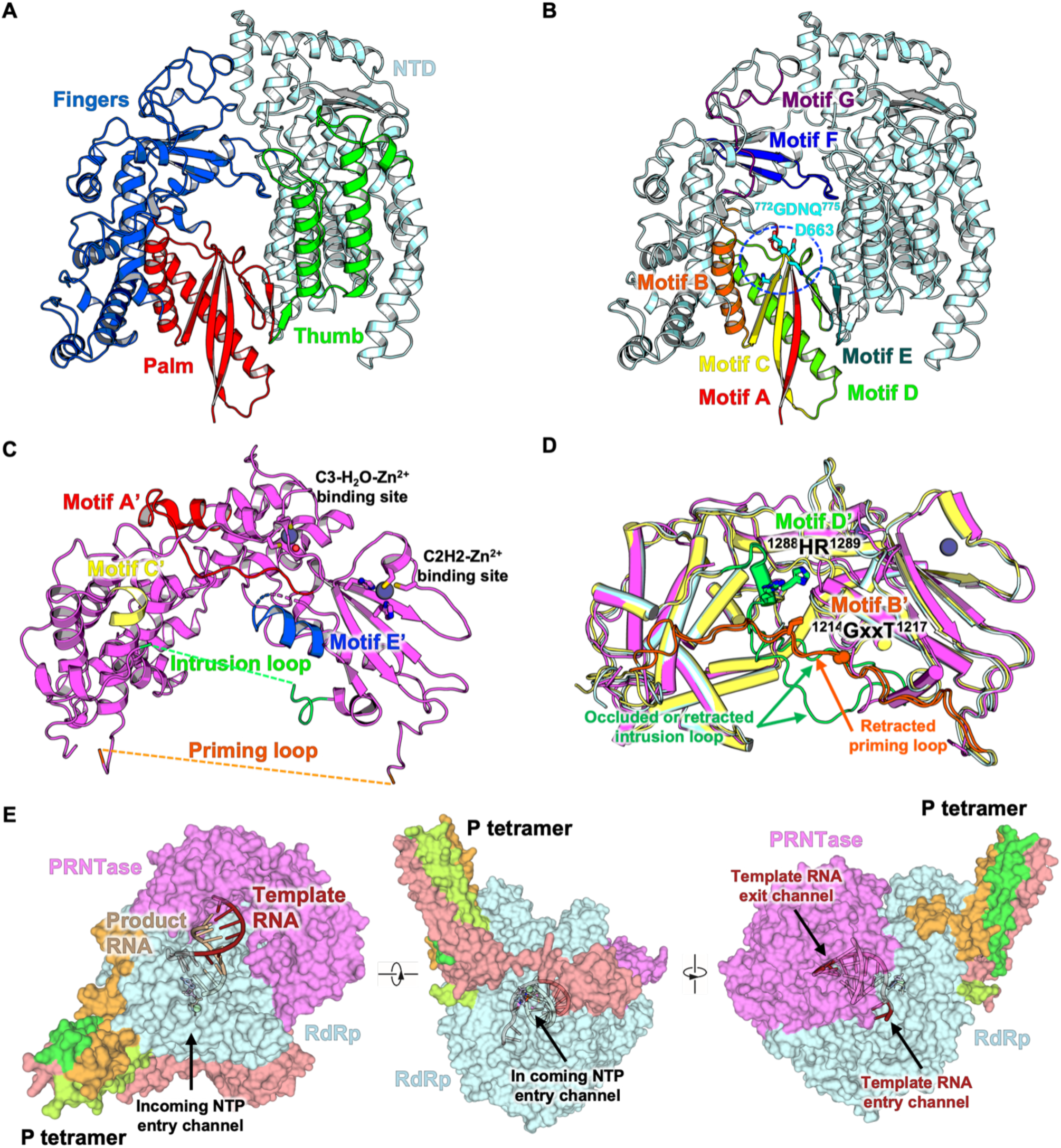
Structural features of the RdRp and PRNTase domains in the MeV apo L–P complex. **A**, Ribbon diagram of the MeV L RdRp domain, colored by subdomain: N-terminal domain (NTD, light blue), fingers (dark blue), palm (red), and thumb (green). **B**, Localization of the seven conserved motifs (A–G) within the RdRp domain. Each motif is color-coded as follows: A (red), B (orange), C (yellow), D (green), E (dark green), F (dark blue), and G (purple). The conserved catalytic residues ^772^GDNQ^775^ in motif C and D663 in motif A are shown as sticks, and the Cα atom of G772 is depicted as a sphere. **C**, Ribbon diagram of the PRNTase domain of MeV L, highlighting three of six visible conserved motifs: A′ (red), C′ (yellow), and D′ (dark blue). The positions of the unresolved priming loop and intrusion loop are marked with orange and green dashed lines, respectively. Two zinc-binding sites are indicated. **D**, Superposition of the experimentally determined PRNTase domain with the two AlphaFold3-predicted MeV L models. Missing regions in the experimented model, including motif B’ (^1288^HR^1289^) and motif D’ (^1214^GxxT^1217^), are highlighted in orange and green, respectively, in the predicted models. The predicted occluded or retracted conformation of intrusion loop and the retracted priming loop are also indicated. **E**, Orthogonal views of the MeV L–P surface representation, accommodated with a model of a dsRNA and an incoming nucleotide. The dsRNA model was derived by superimposing the polymerase core with that of the dsRNA-bound NiV L–P complex (PDB: 9GJU). For clarity, the superimposed NiV structure is not shown.

Like other nsNSVs, the PRNTase domain of MeV L, which is responsible for adding a 5’ cap to nascent viral mRNAs through a unique mechanism, comprises five conserved sequence motifs (motif A’–E’)^5^. In our L–P and LPC structures, only three of these motifs are resolved (Fig. 2C). Two flexible loops, including the priming loop (residue 1208–1231) and the intrusion loop (residues 1290–1302), extend from the PRNTase domain and reflect the catalytic states of the polymerase. In contrast to structures from PIV5, HPIV3, Newcastle disease virus (NDV), and Mumps virus (MuV), where the priming loop is retracted and the intrusion loop projects into the catalytic cavity^59, 60, 61, 62^, both loops are unresolved in the MeV L protein within both the apo L–P and LPC complexes (Fig. 2D). This configuration closely resembles that observed in L structures from RSV, HMPV, NiV, and the state 1 conformation of EBOV^63, 64, 65, 66, 67^. Additionally, the supporting helix from the palm subdomain, which is essential for RNA synthesis, is completely absent in our structure (Fig. 2A), leaving the central cavity accessible for accommodating transcribing RNA. Together, these features indicate that the MeV L–P complex adopts an elongation-state conformation during replication or transcription following cap addition.

The RdRp domain interfaces extensively with the PRNTase domain through contributions from the fingers and thumb subdomains, as well as the N-terminal domain (NTD) of RdRp. This three-dimensional arrangement is essential for maintaining the structural integrity and enzymatic activity of the L protein. When the palm subdomain of the MeV L protein is aligned with the RNA-bound structure of the NiV polymerase, the template RNA entry and exit channels, along with the trajectory of the nascent RNA within the central cavity, are clearly delineated (Figure 2E).

### Bipartite interactions between L and P protein and structural flexibility of the P tetramer

The P protein functions as an essential chaperone of the MeV L protein and plays a critical role in supporting the RNA synthesis^68^. Unlike VSV and Rabies viruses (RABV), whose P proteins form dimers^69, 70^, the MeV P protein assembles into a tetramer through the central coiled-coil domain, a feature shared with the polymerase complexes of RSV, HMPV, EBOV, MARV, HPIV3, PIV5, NDV, MuV and NiV^59, 60, 61, 62, 63, 64, 65, 66, 67, 71^. In all three cryo-EM structures we resolved, each P protomer in the tetramer adopts a different conformation, particularly in the regions following the coiled-coil helices (Fig. 3A). Three of the four P protomers engage with the L protein, primarily contacting the NTD and fingers subdomain of the RdRp via extensive polar interactions across a large interface of ∼3355 Å^2^ (Fig. 3B). Particularly, the disordered segments immediately following the coiled-coil domains of P-1 and P-4 form a short β-sheet with the fingers subdomain, effectively anchoring the P tetramer on the L protein (Fig. 3C and D). P-3 engages the fingers subdomain of L through a combination of polar and hydrophobic interactions, whereas P-2 does not make direct contact with L (Fig. 3E). Additionally, the three-helical bundle of XD domain of P-1 sticks to the NTD via a combination of polar and hydrophobic interactions, further stabilizing the complex (Fig. 3F). In contrast to RSV and HMPV, where the flexible linker of P wraps around the palm to form the NTP entry channel^63, 64^, the corresponding region in MeV P protein loosely traverses the palm subdomain without tight association, likely contributing to its weaker density due to reduced structural stabilization by L (Fig. 3C). Overall, the L protein possesses bipartite binding interfaces that tightly coordinate the P tetramer. In accordance with previous biochemical findings, the N-terminal region of L (residues 1–408) is responsible for P binding and is essential for the RNA synthesis activity of the polymerase^72, 73^.

**Fig. 3.**
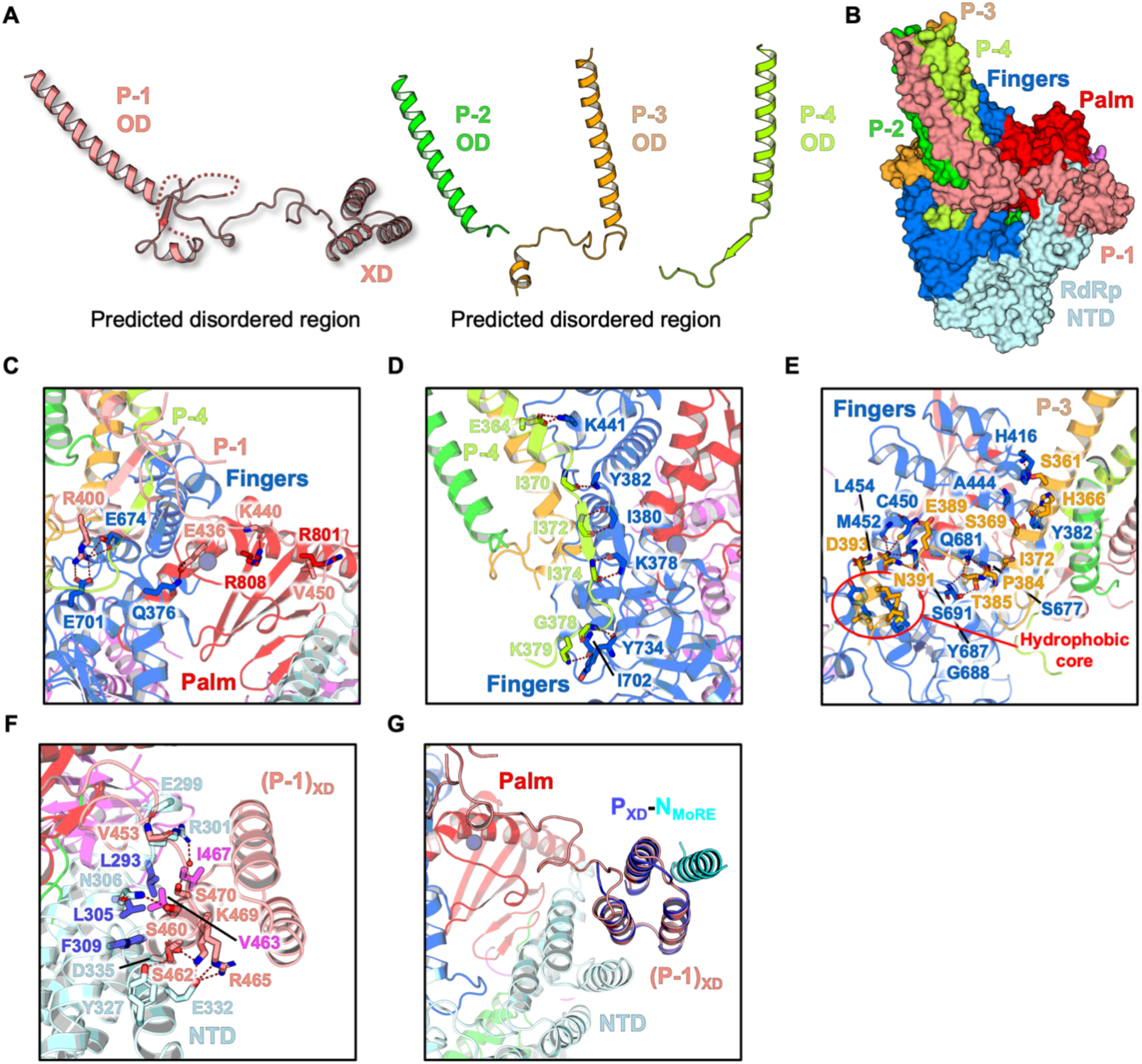
Detailed interaction between MeV L and P proteins. **A**, Structural representations of the four P protomers within the tetramer, highlighting their distinct conformations. The domains of P protein are indicated, including the oligomerization domain (OD), predicted intrinsically disordered regions, and the C-terminal X domain (XD). **B**, Surface representation of the interaction interface between the MeV L and the P tetramer. The subdomains of the RdRp are colored as follows: NTD (light blue), fingers (dark blue) and palm (red). P protomers are colored as in Fig.1A. **C–F**, Close-up views of the specific interaction interfaces between each P protomer and the L protein, illustrating both hydrophobic and polar contacts. **G**, Structural superposition of the crystal structure of the MeV P_XD_–N_MoRE_ complex (PDB: 1T6O) onto the XD region of the P-1 protomer observed in the apo L–P complex.

The XD domain of P molecule was shown to play a key role in replication by associating with the MoRE motif located at the C terminus of N protein. The crystal structure of MeV P_XD_–N_MoRE_ complex could be superimposed well with the P_XD_ portion in our structure (Fig. 3G), indicating that N_MoRE_ engagement does not interfere with the association between P_XD_ and L, and there is no obvious structural rearrangement in the P_XD_ as well. The association between P_XD_ and L is mediated mainly by polar interactions in the solvent exposed region as well as hydrophobic interactions in the buried area, in which residue V463 in the α1 of the P_XD_ plays a key role in mediating the formation of the hydrophobic center in the interface (Fig. 3F) and mutation of this residue impairs the interaction with L protein^72^.

### Structural features of the LPC complex

The C protein has been shown to play a regulatory role in MeV RNA synthesis. In the 2.7 Å-resolution structure of LPC complex, two C molecules are clearly resolved (Fig. 4A). Notably, the non-core region of the L protein containing the CD, MTase and CTD, is observed to fold back onto the PRNTase domain and interacts with one of the bound C proteins (Fig. 4A). Each C protein adopts a compact globular fold composed of six α-helices: one long N-terminal helix followed by five shorter helices arranged nearly perpendicular to each other (Fig. 4B). This fold is structurally distinct from the C proteins of SeV (*Respirovirus*) and Tupaia paramyxovirus (TupV, *Narmovirus*)^74, 75, 76^ (Supplementary Fig. 7A–C). In addition, both C molecules lack visible density for the N-terminal ∼50 residues preceding the first long helix, as well as the linker between the first two helices, indicating these regions are flexible or intrinsically disordered (Fig. 4B). Additionally, weak densities corresponding to the C-terminal tails of both C protomers are present, but insufficient for reliable model building.

**Fig. 4.**
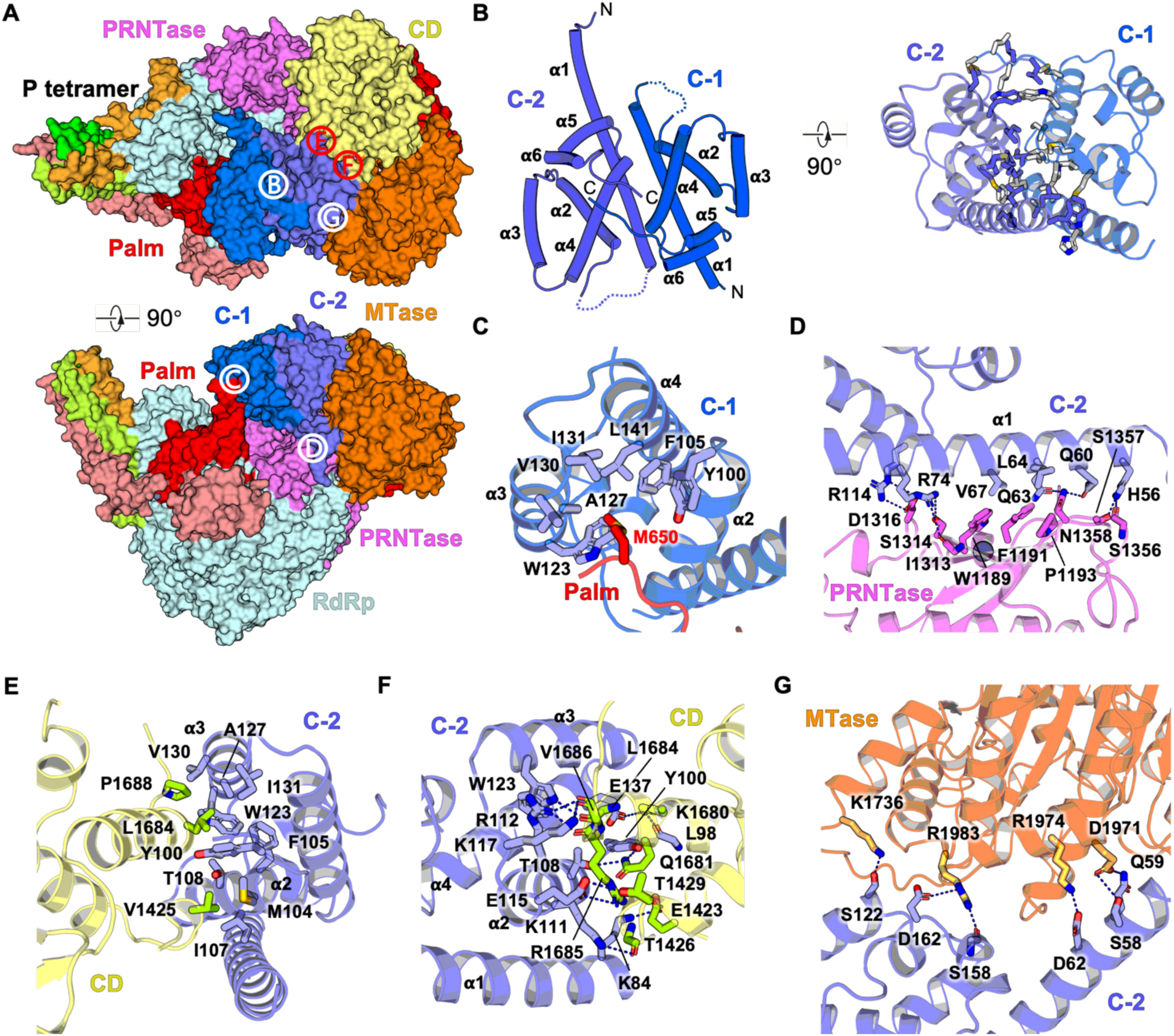
Structural features of the MeV LPC complex. **A**, Surface representation of the interaction interface between the MeV L and the C protein dimer. The subdomains of RdRp and the bound P protomers are colored as in Fig. 3B. The PRNTase domain and the accessory domains of L protein, including the CD, MTase and CTD, are colored as in Fig. 1B. Interaction interfaces between the C protein dimer, as well as those between C protomers and L protein domains, are labeled as **B** to **G**. **B**, Overview of the C protein dimer shown in cartoon (left) and ribbon (right) representations. Residues mediating hydrophobic interactions are displayed as sticks, colored slate and gray. **C–G**, Close-up views of the distinct interaction interfaces between the C protomers and individual domains or subdomains of the L protein, highlighting both hydrophobic and polar interactions that stabilize the complex.

In the LPC structure, the C proteins form a symmetric dimer, which is stabilized primarily by hydrophobic interactions among α-helix 1, 2 and 4 (Fig. 4B), with buried interface area ∼1575 Å^2^. This mode of dimerization is distinct from that observed in the crystal lattice of the TupV C protein (Supplementary Fig. 7D and E), which exists as a monomer in solution, similar to the C proteins of SeV and HPIV1^74, 75, 77^. AlphaFold3-predicted C protein dimer structures from CDV and RPV suggest that these proteins are likely to assemble and function in a manner similar to the MeV C protein (Supplementary Fig. 7F and G), despite their low primary sequence conservation across the morbillivirus genus^24^.

### Detailed interactions between C protein dimer and L

In addition to forming a dimer, both C monomers associate with the polymerase protein through different interaction interfaces in the LPC structure. Residue M650 from the loop prior to the β-strand of motif A in the palm subdomain inserts into the hydrophobic core formed by the α-helices 2, 3 and 4 of one C protomer and plays a role in supporting this C protomer to attach to the RdRp domain (Fig. 4C). The corresponding residue in the closely related CDV and RDV polymerase is a valine, which could also perform the same function as in MeV L. Besides, the other C monomer is located above the PRNTase domain and simultaneously associates with the CD as well as the MTase domain mainly through polar and hydrophobic interactions (Fig. 4D–G), with totally buried interface area ∼1850 Å^2^. The α-helix 1 is the main contacting site for the C protomer to interact with the PRNTase domain and the other two surfaces of the globular C protomer associates with the CD and the MTase domain, respectively.

### Conformational rearrangement of C terminal regions of L upon C dimer engagement

To date, the RNA polymerase structures of nsNSVs that contain fully visible accessory domains include those of VSV and RABV in the rhabdovirus family, as well as HPIV3, PIV5, NDV, and MuV in the paramyxovirus family^59, 60, 61, 62, 78, 79^. Interestingly, unlike other paramyxoviruses, the accessory domains of MeV do not appear to be firmly associated with the core region under either physiological or slightly high salt conditions, indicating a weak interaction between them (Supplementary Fig. 8). However, upon the association of the C dimer, the MTase and CTD become stabilized and adhere to the PRNTase domain through distinct interaction interfaces compared to other viruses (Supplementary Fig. 9A and B), whereas the CD has few contact with the PRNTase domain.

Among RNA polymerase structures with visible accessory domains, the spatial arrangements of these domains are highly different and adopt distinct conformations, reflecting various functional states during the certain stages of replication and transcription (Fig. 5). In the VSV and RABV polymerase structures (PDB: 5A22, 6U1X and 6UEB) that assume in the pre-initiation state, the deep insertion of the priming loop from the PRNTase domain into the catalytic cavity, the supporting helix in the palm subdomain, and the entire CD located above the central cavity obstruct the exit channel of the nascent RNA transcript, so that the withdrawal of the priming loop, displacement of the supporting helix, as well as reorganization of the accessory domains are required for the growing RNA product^78, 79^ (Fig. 5A and B). In the PIV5 polymerase structure (PDB: 6V85) that adopt in the post-initiation state, despite the reorganization of the accessory domains compared to the VSV and RABV polymerases, the intrusion loop from the PRNTase domain, the supporting helix, as well as the CD prevent the nascent RNA to extend out from the central cavity^59^ (Fig. 5G). In the HPIV3 and NiV polymerase structure (PDB: 8KDB and 9GJU), in which the priming loop has been retracted, and the intrusion loop is partially missing but the trajectory still projects into the catalytic cavity, the supporting helix and the CD still block the exit tunnel for the nascent RNA product^61^ (Fig. 5C and D). The states of the priming loops, the intrusion loops and the supporting helices in both the NDV and MuV polymerase structures (PDB: 7YOU and 8IZl) are quite similar to those in the HPIV3 polymerase structure (PDB: 8KDB), however, the CDs are already elevated and apart from the RdRp and PRNTase domains compared to the HPIV3 polymerase, partially exposing the potential product RNA exit channel^60, 62^ (Fig. 5E and F).

**Fig. 5.**
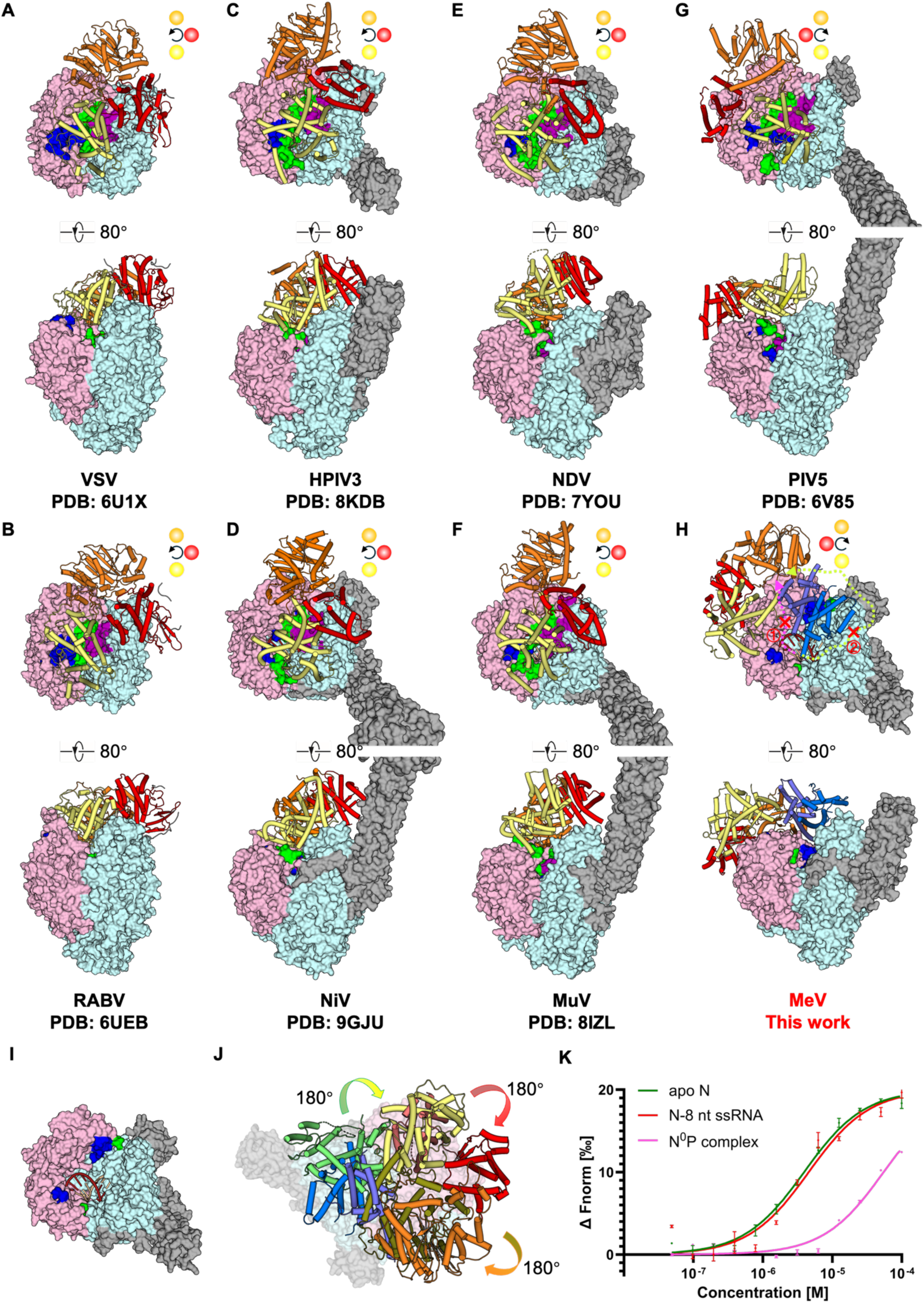
Comparison of the catalytic states among various nsNSV polymerase structures. The cores of the polymerases, including the RdRp and PRNTase domains, are shown as surface representations, colored light blue and pink, respectively. The priming loops, intrusion loops and supporting helices are displayed as surfaces in green, dark blue and deep magenta, respectively. The accessory domains of the L protein, comprising the CD, MTase and CTD, are shown as cartoons in yellow, orange and red, respectively. P protein tetramers are displayed as gray surfaces. The spatial positions of the accessory domains are further indicated by colored circles in top-down views: yellow for CD, orange for MTase, and red for CTD. Clockwise and counterclockwise arrows denote the spatial rearrangements of these domains. **A** and **B**, Structural features of the rhabdovirus polymerases VSV (PDB: 6U1X) and RABV (PDB: 6UEB) in the pre-initiation state, showing obstructing priming and intrusion loops and a closed accessory domain arrangement. **C** and **D**, Structures of HPIV3 (PDB: 8KDB) and NiV (PDB: 9GJU) polymerases, showing conformations in which the intrusion loop, supporting helix and CD occlude the product RNA exit tunnel. **E** and **F**, NDV (PDB: 7YOU) and MuV (PDB: 8IZL) polymerases in a conformation with a partially exposed product RNA exit channel. **G**, Structure of PIV5 (PDB: 6V85) polymerase in the post-initiation state, showing further rearrangement of accessory domains. **H** and **I**, Spatial configuration of accessory domains (CD, MTase, CTD) in the MeV LPC complex (**H**) and structural view of the core region (**I**) of LPC complex. The priming loop, intrusion loop, and supporting helix are largely unresolved (**I**) and the product RNA exit channel is completely open (**H**). The RNA duplex is modeled as in Fig. 2E. Two potential exit routes for the product RNA toward the MTase active site are indicated by dashed curves: route 1 in magenta and route 2 in green. **J**, Conformational comparisons of CD, MTase and CTD between PIV5 and MeV polymerase. PIV5 domains are shown in green (CD), brown (MTase), and salmon (CTD), with arrows indicating domain rotation angles. For clarity, the superimposed PIV5 core structure is omitted. **K**, MST analysis of full-length C protein binding to different forms of N proteins, including the monomeric apo N protein, N protein bound to 8 nt ssRNA, and the N^0^P complex. Data represent means ± S.D. from n = 3 biologically independent replicates.

The state of our MeV LPC complex structure is quite different from all the previously determined structures (Fig. 5H). Due to missing the complete the priming loop, the intrusion loop as well as the supporting helix, the central cavity is largely vacant (Fig. 5H). In addition, as a result of the association of the C protein dimer, the CD-MTase-CTD module is completely displaced from the above of the central cavity and only attaches to the PRNTase domain of the L protein, fully exposing the product RNA exit channel (Fig. 5I). The domain organization of the accessory domains in the MeV LPC complex is similar to that in the PIV5 polymerase structure (PDB: 6V85), in which the CD-MTase-CTD are arranged in an anticlockwise orientation from the aerial view, whereas in the polymerase structures of several other viruses mentioned above, these domains are arranged in a clockwise direction (Fig. 5H). Through superimposing the core of the PIV5 polymerase (PDB: 6V85) onto that of the LPC structure, it appears that the C protein dimer causes a dramatic rotation and shift of the entire CD-MTase-CTD module, in which the CD flips nearly 180°, the MTase domain rotates about 86° and the CTD about 76° (Fig. 5J). The elongating RNA transcript emerging from the product exit channel seems unable to travel around the CD-CTD-MTase route 1 to reach the active site of the MTase domain like in the polymerase structure of NDV or MuV (Fig. 5H). In addition, the product RNA also cannot pass through the C dimer-MTase route 2 or directly penetrate through the interior tunnel to access the catalytic sites of the MTase domain (Fig. 5H), due to the interaction between C-2 and CD (Fig. 4E and F). Thus, the state we presented here is unlikely to be the capping or methylation reaction stage during the transcription process.

In order to investigate the role and potential interaction partners of the C protein in the RNA synthesis process, we performed the MST analysis screen using eGFP-tagged C protein as a target. In addition to the MeV LP complex, we also found that the C protein is able to interact with the N protein directly. The MST results showed that the affinity of the C protein between the monomeric apo N and the N–8 nt ssRNA complex is about 3.92 ± 0.50 μM and 4.57 ± 1.46 μM, respectively, while the affinity between the C protein to N^0^P complex is dramatically decreased (Fig. 5K). Thus, the interaction between N and C uncovers another critical function of C that not only opens up the nascent RNA exit channel but possibly plays a role in recruiting the monomeric N protein to the site where the RNA product emerges from the catalytic cavity, which is poised to encapsidate the nascent RNA. Since only the newly-synthesized viral genome and antigenome are encapsidated by N proteins to form the nucleocapsid during the RNA synthesis process, our LPC structure determined here reflects a bona fide elongation state during viral replication, and the LPC complex could thus be regarded as a replicase whereas the L–P complex is just a transcriptase, which also indicates that the non-structure protein C regulates the switch from transcription to replication through association with N protein.

### Structural basis of ERDRP-0519 recognition by the MeV L protein

A series of non-nucleotide small-molecule inhibitors targeting the morbillivirus polymerase complex have been developed, including 16677, AS-136a and ERDRP-0159 (Supplementary Fig. 10 A–C), all of which revealed nanomolar potency. Surprisingly, contrary to previous photo-crosslinking and *in silico* docking studies suggesting that the binding pocket of ERDRP-0519 is located in the PRNTase domain of L^55^, our 3.1 Å-resolution cryo-EM map and structure of the L–P complex bound to ERDRP-0519 (hereafter referred to as the LPE complex) identifies extra density positioned in the cavity surrounded by the fingers and palm subdomain of the RdRp (primarily involving motifs A–D), in the vicinity of the ^772^GDNQ^776^ catalytic sites (Fig. 1D, Fig. 6A and B). This density is absent in the corresponding region of both apo L–P and LPC maps. Notably, no additional density is detected in the PRNTase domain of the LPE map, even at near-atomic resolution, confirming that ERDRP-0519 binds within the RdRp domain rather than the previously proposed PRNTase site. The inhibitor occupies a buried pocket (∼500 Å^2^ interface area) deep within the RdRp catalytic cavity. Furthermore, ERDRP-0519 and its first-generation analogues fit the density unambiguously when modeled with appropriate geometry restraints (Supplementary Fig. 10 D–F).

**Fig. 6.**
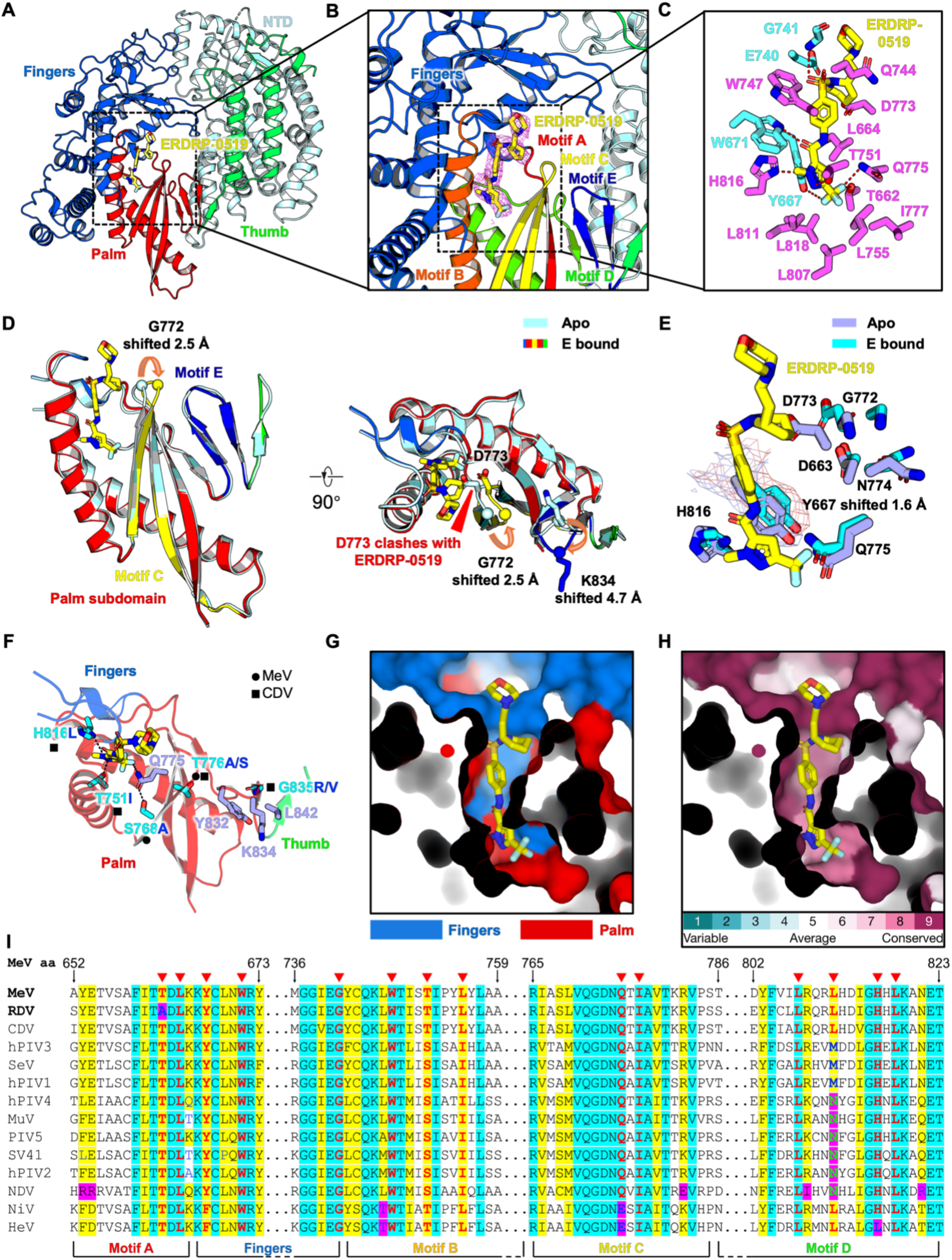
Molecular mechanism of ERDRP-0519 recognition and inhibition of MeV RNA polymerase. **A**, Spatial localization of ERDRP-0519 within the RdRp domain of the MeV L protein. Subdomains are colored as in Fig. 2A. ERDRP-0519 is shown as yellow sticks. **B**, Close-up view of the ERDRP-0519 binding pocket formed by the palm and fingers subdomains. Motifs A to E within the palm subdomain are colored red (A), orange (B), yellow (C), green (D), and dark blue (E). The cryo-EM density for ERDRP-0519 is shown as a magenta mesh. **C**, Detailed interactions between ERDRP-0519 and the RdRp. Key residues from the palm and fingers subdomains are shown as magenta and cyan sticks, respectively. Potential polar interactions are indicated by red dashed lines. **D**, Structural superposition of the ERDRP-0519 binding pocket in the apo and ERDRP-0519-bound MeV L–P complexes. Clashes between ERDRP-0519 and the apo conformation are indicated, along with conformational rearrangements in motifs C (yellow) and E (dark blue) upon inhibitor binding (highlighted by red arrows). The apo structure is shown in light blue; the inhibitor-bound structure is colored as in panels **A** and **B**. **E**, Conformational differences in main and side chains of key binding residues between the apo and ERDRP-0519-bound structures. Residues from the apo structure are shown as light slate sticks, while those from the inhibitor-bound structure are shown as cyan sticks. Cryo-EM densities for residue Y667 are displayed as meshes in light blue (apo) and salmon (bound). **F**, Mapping of resistance mutations. Escape mutations identified in MeV (small black dots) and CDV (black squares) polymerases are shown as cyan sticks in the ERDRP-0519 binding pocket. Surrounding residues are shown as light slate sticks. **G** and **H**, Surface representation of the ERDRP-0519 binding pocket. Panel **G** indicates the spatial location of the pocket, while panel **H** displays sequence conservation calculated using the ConSurf server^95^. **I**, Structure-based partial sequence alignment of representative paramyxoviral L proteins, encompassing regions of motif A, the fingers subdomain, and motifs B, C, and D, which contribute to the ERDRP-0519 binding pocket. Residues directly involved in inhibitor binding are shown in bold red font and marked with red triangles above the alignment. Absolutely conserved residues are highlighted in cyan, and relatively conserved residues in yellow. Residues with conservative substitutions are in bold blue, while non-conservative substitutions are in bold green. All residues at or near the binding pocket that differ from MeV are highlighted in magenta.

The LPE structure provides detailed view of how this compound is specifically recognized by the MeV L protein (Fig. 6C). Notably, the N-ethylmorpholinyl group at the position 2 of the piperidine ring extends into the solvent exposed central cavity formed by the RdRp and PRNTase domains, without making direct contact with surrounding residues (Fig. 6C). This hydrophilic group was introduced primarily to enhance aqueous solubility and oral bioavailability, with minimal impact on antiviral potency compared to its lead compound, AS-136a^52^. The piperidine ring is partially buried and stabilized through weak Van der Waals’ interactions with the alkyl chains of surrounding polar residues E740, Q744 and D773 from the catalytic ^772^GDNQ^776^ motif (Fig. 6C). A hydrophobic residue, L664, supports the ring from below, while a hydrogen bond between G741 and the sulfonyl linker further anchors the molecule in place (Fig. 6C). Aromatic residue W747 forms a perpendicular π–π stacking interaction with the central phenyl ring, which is further clamped by L664 via hydrophobic contact from the other side (Fig. 6C). The acylamino linker is additionally stabilized by a hydrogen bond from W671 (Fig. 6C). The most deeply buried portion of the inhibitor, the 1-methyl-3-trifluoromethyl-pyrazole moiety, is inserted into a highly hydrophobic pocket formed by residues L664, Y667, W671, L755, L777, L807, L811, and L818. Within this pocket, Y667 forms another perpendicular π–π interaction with the pyrazole ring (Fig. 6C). Several polar contacts, involving T662, Y667, T751, H816 as well as Q775 from the ^772^GDNQ^776^ motif, interact with nitrogen and fluorine atoms of the pyrazole and its CF₃ substituent, further reinforcing the binding (Fig. 6C). Together, these extensive hydrophobic and polar interactions encase ERDRP-0519 with high specificity, explaining its tight binding to MeV L with a reported affinity ∼80 nM^55^, which is the prerequisite for the antiviral efficacy of the drug *in vivo*.

### Molecular mechanism of MeV RNA synthesis inhibition by ERDRP-0519

We then explored the molecular mechanism by which the non-nucleotide inhibitor ERDRP-0519 blocks the RNA synthesis activity of MeV and CDV polymerase complexes. Structural superposition of the fingers–palm subdomain in the LPE complex with that of the apo L–P structure revealed a remarkable conformational change in the β-hairpin loop of motif C, where the catalytic ^772^GDNQ^776^ motif resides (Fig. 6D). Particularly, the Cα atom of residue G772 moves 2.5 Å away from the inhibitor, and the catalytic residue D773 reorients to the opposite side to avoid the steric clash with the piperidine ring of the inhibitor (Fig. 6D). These shifts propagate to motif E, inducing a rearrangement of its β-hairpin loop and resulting in a 4.7 Å displacement of the Cα atom of K834 at its tip (Fig. 6D). Furthermore, inhibitor binding alters the interaction network between motifs C and E. In the apo state, Q771 in motif C forms a hydrogen bond with the side chain of S833, D773 forms a salt bridge with R544 in motif F, and N774 interacts with the backbone of V831 in motif E (Supplementary Fig. 10G). Upon inhibitor engagement, Q771 instead interacts with the backbone of S833, while S833 forms a new hydrogen bond with G772. Additionally, D773 establishes a novel interaction with K665 in motif A, further disrupting the native catalytic environment (Supplementary Fig. 10G). Similar local conformational changes were observed when the LPC structure was superimposed with the LPE model (Supplementary Fig. 10H), indicating that the inhibitor ERDRP-0519 can also bind to the fingers–palm subdomain of LPC complex and block both transcription and replication.

In addition to the structural rearrangements at the catalytic center, several residues within the ERDRP-0519 binding pocket also experience local conformational adjustments upon inhibitor engagement. Notably, Y667 shifts outward by 1.6 Å to maintain favorable Van der Waals interactions (Fig. 6E). Meanwhile, Q775 and H816 move closer to the compound, adopting new side-chain rotamers to optimize binding contacts (Fig. 6E) These subtle but coordinated changes suggest an induced-fit binding mode rather than a rigid lock-and-key mechanism. Such conformational plasticity in the palm subdomain likely distorts the optimal geometry of the active site, thereby compromising the catalytic competence of the polymerase. Additionally, ERDRP-0519 appears to lock the RdRP complex into an elongation-like conformation, preventing the conformational transitions required for initiation, re-initiation, and termination. This structural rigidity may further impair polymerase turnover during RNA synthesis, offering a mechanistic explanation for the potent antiviral efficacy observed in both cell-based assays and animal studies.

To investigate mechanisms of resistance, we analyzed previously reported escape mutations in the MeV and CDV L proteins that emerged during viral adaptation in the presence of ERDRP-0519 or its precursor compounds^48, 55^. These resistance mutations show high convergence between MeV and CDV, clustering within functionally critical regions of the polymerase (Fig. 6F, Supplementary Fig. 10I). In CDV, the N398D mutation resides in the fingers subdomain, while in MeV, L1170F, R1233Q, and V1239A are found in the PRNTase domain (Supplementary Fig. 10I and J). Additional mutations—T751I, H816L, and G835R/V in CDV, as well as H589Y and T776A/S shared by both viruses—are located in the palm subdomain (Fig. 6F).

Our LPE complex structure provides insight into the resistance mechanism. Residues T751 and H816 form direct polar contacts with the pyrazole moiety of ERDRP-0519, while S768 stabilizes Q775, which in turn hydrogen-bonds with the CF_3_ group of the compound (Fig. 6F). Thus, mutations at these sites are likely to disrupt binding through either steric hindrance or loss of polar interactions. Indeed, the S768A mutation causes over 400-fold decrease in binding affinity and 28-fold increase in EC_50_ in the cell-based assays^55^. Similarly, T776, part of the β-strand in motif C, contributes to local structural integrity^80^. Its mutation (e.g., T776A or T776S) can destabilize the β-sheet, leading to an ∼80-fold reduction in compound sensitivity^55^. Residue G835, located in motif E and surrounded by Y832, K834, and L842 from the thumb subdomain, is particularly sensitive to side-chain volume changes (Fig. 6F). Substitution with bulkier residues like valine or arginine may disrupt local packing, altering the geometry of motifs E and C and diminishing compound efficacy. H589, although not resolved in our elongation-state structure, is predicted to locate in the supporting helix and to hydrogen bond with G840 (Supplementary Fig. 10I), a highly conserved residue connecting the motif E of the palm and the thumb subdomain. The H589Y mutation may weaken this interaction and impact structural communication between palm and thumb subdomains. Outside the immediate binding pocket, N398 in the fingers subdomain (mutated to D in CDV) interacts with the main chain of L431 (Supplementary Fig. 10J). Another three mutants, L1170F, R1233Q and V1239A, located in the PRNTase domain, are distant from the inhibitor-binding cavity and mutations in these sites are unlikely to directly affect the architecture of the palm subdomain but might induce allosteric effects, subtly altering polymerase conformation or enhancing RdRP basal activity, thereby reducing inhibitor susceptibility.

Viruses within the *Paramyxoviridae* family typically exhibit a high degree of structural and sequence conservation, particularly in the L protein (Supplementary Fig. 11). In order to explore whether the non-nucleotide inhibitor ERDRP-0519 could potentially bind and inhibit the polymerases from other paramyxoviruses beyond morbilliviruses, we performed structure-guided sequence alignments based on conserved motifs and residues directly involved in compound binding within the MeV L protein. Both sequence alignment and binding pocket analysis revealed a remarkable conservation of residues forming the inhibitor-binding cavity as well as their immediate neighbors (Fig. 6G–I), suggesting that the antiviral efficacy may extend beyond MeV and CDV to the other genera within the family. However, several site-specific variations were observed. For example, T662 in MeV is replaced by alanine in RDV, potentially disrupting a hydrogen bond with the inhibitor (Fig. 6I). In addition, L811 in MeV is substituted by methionine in HPIV3, SeV and HPIV1, a conservative change that likely preserves hydrophobic interactions (Fig. 6I). Intriguingly, in rubellavirus, L811 is replaced by asparagine (Fig. 6I), which may reduce hydrophobic contact but introduce a new potential hydrogen bond with the CF_3_ group of the compound. Notably, the highly conserved glutamine in the GDNQ motif of nsNSV is substituted by glutamate in NiV and HeV (Fig. 6I), possibly weakening polar interactions with the inhibitor. Additionally, H816 in MeV is replaced by leucine in HeV (Fig. 6I), altering the local electrostatic environment of the pocket. These residue-level differences could modulate the binding affinity of ERDRP-0519 across *Paramyxoviridae* polymerases and highlight the potential to expand this pan-morbillivirus inhibitor into a broader-spectrum pan-paramyxovirus antiviral agent.

### Mechanistic insights of ERDRP-0519 engagement with NiV polymerase

Given that several amino acid substitutions are present in the putative ERDRP-0519 binding site of NiV and HeV polymerases, we aimed to investigate whether ERDRP-0519 can still engage these viral polymerases and retain its inhibitory effect. To this end, we determined the cryo-EM structure of the NiV polymerase bound to this compound at 2.8 Å resolution (Fig. 7A and B, Supplementary Fig. 12 and 13). The resulting map revealed an additional density located within a narrow cavity formed between the fingers and palm subdomains of the L protein (Fig. 7A), consistent with the binding of ERDRP-0519 (Fig. 7B–D). Overall, the compound adopts a binding mode in the NiV polymerase that is highly similar to that observed in the MeV polymerase, although subtle differences in residue interactions are evident (Fig. 7E).

**Fig. 7.**
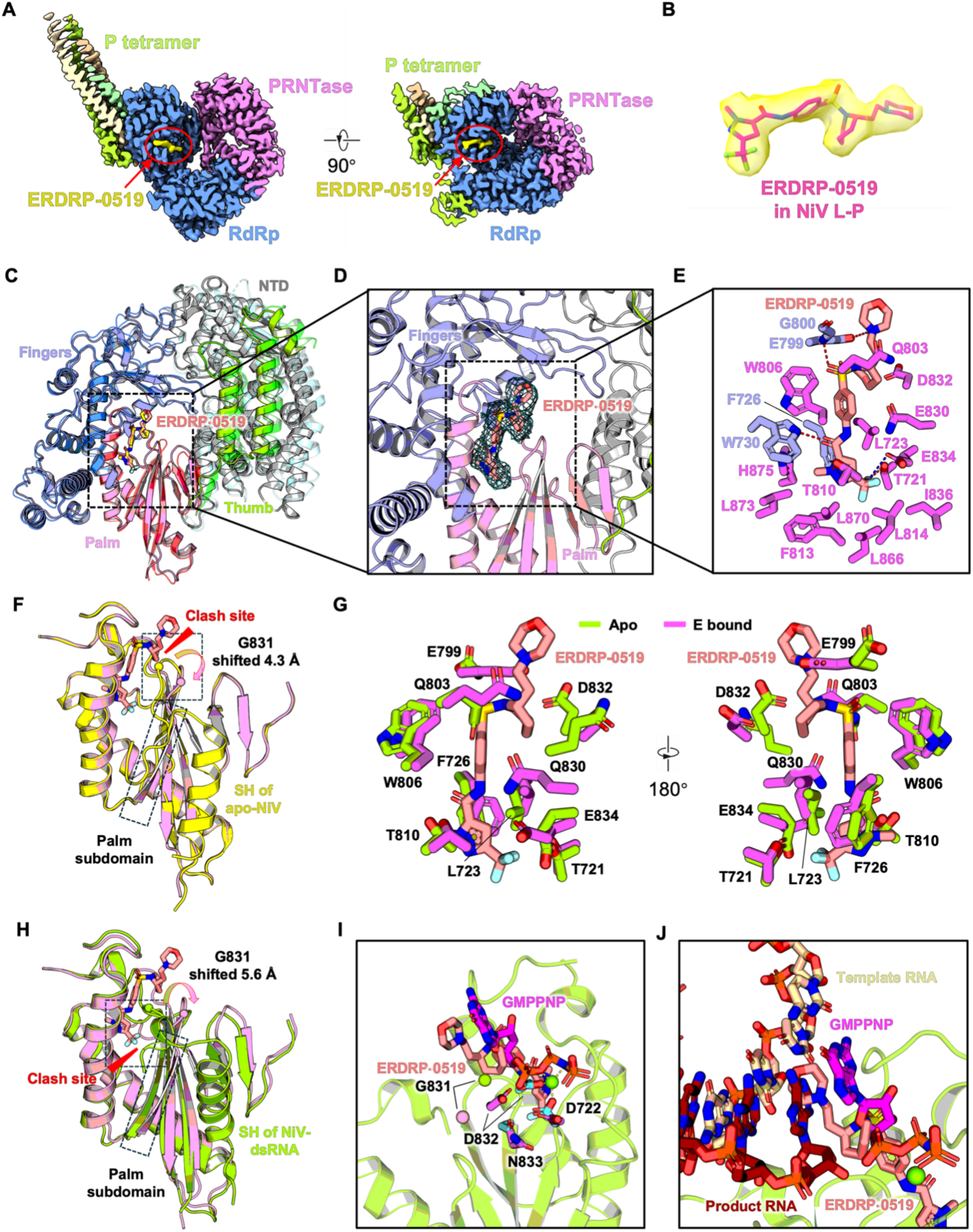
Mechanistic insights into ERDRP-0519 binding to and the inhibition of NiV polymerase. **A**, Cross-sectional view of the cryo-EM map of the NiV L–P complex bound to ERDRP-0519. The RdRp and PRNTase domains are colored in blue and pink, respectively. The four P protomers are shown in light orange, wheat, limon and lime. The density corresponding to ERDRP-0519 is highlighted in yellow within the RdRp domain. **B**, Zoomed-in view of the ERDRP-0519 density with the fitted atomic model overlaid. **C**, Comparison of the spatial localization of ERDRP-0519 within the RdRp domains of the MeV and NiV L proteins. Subdomains of MeV RdRp are colored as in Fig. 2A. The NiV RdRp subdomains are colored as: NTD (gray), fingers (light slate), palm (light pink) and thumb (limon). ERDRP-0519 is shown as yellow (MeV) and salmon (NiV) sticks. **D**, Close-up view of the ERDRP-0519 binding pocket formed by the palm and fingers subdomains of NiV RdRp. The cryo-EM density for ERDRP-0519 is shown as a dark blue mesh. **E**, Detailed interactions between ERDRP-0519 and the RdRp. Key residues from the palm and fingers subdomains are shown as magenta and light slate sticks, respectively. Potential polar interactions are indicated by red and blue dashed lines. **F**, Superposition of the ERDRP-0519 binding pocket in the apo (PDB: 9BDQ) and inhibitor-bound NiV RdRp complexes. Clashes between ERDRP-0519 and the apo conformation are indicated by red arrows. Conformational changes within the palm subdomain upon inhibitor binding are highlighted. The apo structure is shown in yellow; the inhibitor-bound structure is colored in light pink. **G**, Conformational changes in side chains of key binding residues between the apo (PDB: 9BDQ) and ERDRP-0519-bound structures. Residues from the apo structure are shown as limon sticks, and those from the inhibitor-bound structure as magenta sticks. **H**, Superposition of the ERDRP-0519 binding pocket in the dsRNA-bound (PDB: 9GJU) and inhibitor-bound NiV L–P complexes. Clashes between ERDRP-0519 and the dsRNA-bound conformation are indicated by red arrows. Conformational changes within the palm subdomain upon inhibitor binding are highlighted. The dsRNA-bound structure is shown in limon; the inhibitor-bound structure is colored in light pink. **I**, Steric hindrance between ERDRP-0519 and the bound nucleotide analogue GMPPNP (magenta sticks) from the dsRNA-bound NiV L–P complex structure (PDB: 9GJU). Catalytic residues from the dsRNA-bound (PDB: 9GJU) structure are shown as cyan sticks, with the Cα atom of G831 as a limon sphere and Mg^2+^ as green sphere. The same residues in the inhibitor-bound structure are shown as magenta sticks, with the Cα of G831 as a light pink sphere. The inhibitor-bound structure is omitted for clarity. **J**, Steric hindrance between ERDRP-0519 and the template and product RNA strands (shown in wheat and firebrick sticks, respectively) from the dsRNA-bound NiV L–P complex structure (PDB: 9GJU). The inhibitor-bound structure is hidden for clarity.

Specifically, the ethylmorpholinyl group at position 2 of the piperidine ring is fully solvent-exposed, with only the carboxyl group of residue E799 forming a weak salt bridge with the protonated tertiary amine of the morpholine ring at neutral pH (Fig. 7E). The piperidine ring is surrounded by the side chains of E799, Q803, Q830 and D832, primarily through Van der Waals’ interactions (Fig. 7E). As seen in the MeV polymerase, the main chain nitrogen atom of conserved G800 forms a hydrogen bond with the connecting sulfonyl group (Fig. 7E). The central phenyl ring is clamped by residues W806 and L723 from the top and bottom via π–π stacking and hydrophobic interactions, respectively (Fig. 7E). The connecting acylamino bond also forms a hydrogen bond with W730 (Fig. 7E). However, unlike in the MeV structure, the equivalent residue, E834 from the ^831^GDNE^834^ catalytic motif does not generate a hydrogen bond with the compound (Fig. 7E). The 1-methyl-3-trifluoromethyl-pyrazole moiety is deeply embedded in a hydrophobic core formed by residues L723, F726, W730, F813, L814, I836, L866, L870, L873 and H875 (Fig. 7E). Additionally, T810 form a hydrogen bond with N2 atom in the pyrazole ring, while T721 interacts with one of the fluorine atoms in the CF_3_ group (Fig. 7E). E834 establishes an unusual charge interaction with the carbon atom of the CF_3_ group, likely driven by the strong electronegativity of the fluorine atoms (Fig. 7E). Together, these polar interactions within the hydrophobic cavity further enhanced the binding affinity of ERDRP-0519 to the NiV polymerase.

Similar to the conformational changes observed in the MeV polymerase upon ERDRP-0519 binding, the NiV polymerase also appears to undergo comparable structural rearrangements in response to the compound (Fig. 7F, Supplementary Fig. 14A and B). The most striking conformational change is the structural remodeling of the motif C within the palm subdomain, particularly the flipping out of the catalytic ^831^GDNE^834^ motif and a noticeable bulging of the main chain in one of the β-strands (Fig. 7F, Supplementary Fig. 14B and C). Additionally, motif A and motif D exhibit slight outward shifts upon compound engagement. These collective movements result in the complete disappearance of the supporting helix, which is likely repelled during conformational rearrangement. Furthermore, residues such as L723, F726 and W806, which are involved in compound binding, also experience corresponding adjustments to facilitate the interaction (Fig. 7G).

To investigate whether and how ERDRP-0519 binding affects RNA synthesis by the NiV polymerase, we superimposed the ERDRP-0519-bound NiV polymerase structure with the RNA- and incoming nucleotide-bound NiV polymerase structure in the early elongation state^67^ (PDB: 9GJU), using the fingers-palm subdomains for structural alignment (Fig. 7H). Notably, in the RNA- and NTP-bound structure (PDB: 9GJU), the catalytic ^831^GDNE^834^ motif flips towards the fingers subdomain, positioning the carboxyl groups of residues D722 from motif A, D832, the carbonyl group of L732, along with the phosphodiester moiety of the incoming GTP analogue, to chelate a Mg^2+^ ion^67^. In contrast, in the ERDRP-bound structure, the ^831^GDNE^834^ motif instead flips toward motif E (Fig. 7H, Supplementary Fig. 14D–F), and the catalytic residues are misaligned, rendering them unable to participate in such coordination (Fig. 7I, Supplementary Fig. 14E). Strikingly, the piperidine ring of ERDRP-0519 directly clashes with the ribose moiety of the GTP analogue (Fig. 7I), while the ethylmorpholinyl group that previously considered to play a negligible role in RNA synthesis inhibition introduces steric hindrance with both the template and product RNA bases (Fig. 7J). These findings reveal that the compound inhibits RNA synthesis not only by disrupting the active-site conformation but also by blocking the binding of the incoming nucleotide and interfering with the positioning of the template and product RNA strands, mechanisms that had not been fully appreciated before.

### Broad-Spectrum targeting of paramyxoviral polymerases by ERDRP-0519

To gain deeper insights into the interaction mechanisms of ERDRP-0519 at its binding site, we conducted a series of molecular dynamics (MD) simulations using the MeV L/ERDRP-0519 and NiV L /ERDRP-0519 cryo-EM structures. Three independent 200-ns simulations were performed for each system. In the MeV simulations, ERDRP-0519 forms stable hydrogen bonds with residues T662, Y667, T751, H816, W671, and Q775, and π–π stacking interactions with Y667 and W747 (Fig. 8A). In the NiV simulations, ERDRP-0519 exhibited a comparable interaction network, indicating a conserved mode of binding across both viruses (Fig. 8B). Critically, Notably, the ligand maintained a binding pose highly consistent with the initial cryo-EM model throughout the trajectories, validating the observed interaction geometry.

**Fig. 8.**
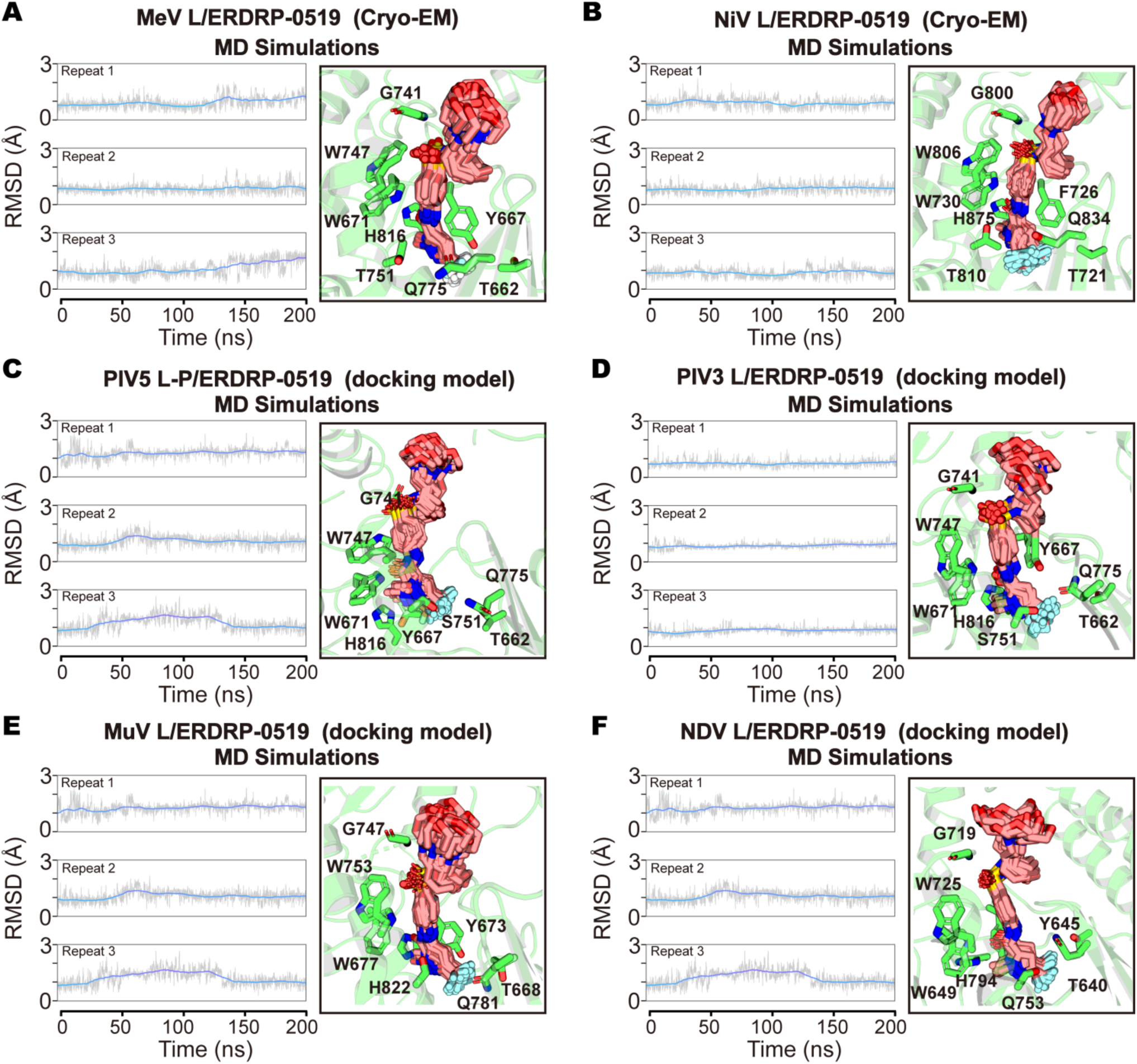
Dynamics of ERDRP-0519 in representative paramyxoviral polymerase complexes. MD simulations and docking analyses of ERDRP-0519 bound to the L–P complex structures of MeV (**A**, this study), NiV (**B**, this study), PIV5 (**C**, docking model), HPIV3 (**D**, docking model), MuV (**E**, docking model), and NDV (**F**, docking model). Three independent 200-ns simulations were conducted for each system. **Left panels:** Root-mean-square deviation (RMSD) plots of ERDRP-0519 over time across the three replicates, illustrating the stability of the compound within the binding pocket. **Right panels:** Representative structural snapshots extracted at 20-ns intervals throughout the simulations. Key conserved residues (e.g., G741, W747) mediating ERDRP-0519 binding are shown as green sticks, and are highly conserved across polymerases. ERDRP-0519 is depicted in salmon, highlighting its consistent binding pose across different *Paramyxoviridae* RdRp domains.

Beyond its established efficacy against MeV polymerase complex, whether ERDRP-0519 exhibits broad-spectrum inhibitory activity against other *Paramyxoviridae* members such as PIV5, HPIV3, MuV, and NDV remains incompletely understood. To explore the potential for cross-viral inhibition, we performed molecular docking and MD simulations using the L proteins of these viruses. Docking results revealed that ERDRP-0519 adopts highly similar binding poses across these polymerases (Supplementary Fig. 15A– D), and MD simulations demonstrated that the compound remained stably bound within the RdRp domain in each case. In the PIV5 system, residues G741, T662, Y667, S751, H816, and Q775 formed hydrogen bonds with ERDRP-0519, while W747 engaged in a perpendicular π–π interaction with its central phenyl ring (Fig. 8C). A nearly identical interaction pattern was observed in HPIV3, involving the same set of residues (Fig. 8D). In the MuV system, ERDRP-0519 formed hydrogen bonds with G747, T668, Y673, H822, W677, and Q781, and π–π stacking interactions with W753 and H822 (Fig. 8E). Similarly, in the NDV polymerase, the inhibitor engaged G719, T640, Y645, H794, W671, and Q753 through hydrogen bonds and formed a π–π stacking interaction with W725 (Fig. 8F). Notably, interactions involving G741 (or its equivalent), W671, and W747 are conserved across these systems, including MeV (G741, W671, W747), PIV5 (G741, W671, W747), HPIV3 (G741, W671, W747), MuV (G747, W677, W753), and NiV (G719, W649, W725). These results suggest that ERDRP-0519 engages a structurally conserved allosteric pocket in the RdRp domain, supported by a recurrent network of conserved residues. While further experimental validation is needed, this structural conservation may underlie the potential of ERDRP-0519 as a broad-spectrum inhibitor targeting multiple *Paramyxoviridae* polymerases.

To investigate why ERDRP-0519 fails to inhibit RNA synthesis in RSV, we performed structural comparisons by superimposing the fingers–palm subdomain of the MeV LPE complex onto that of the RSV polymerase in the elongation state. The analysis revealed several steric clashes between the compound and key residues in RSV (Supplementary Fig. 16A and B). Notably, residues F704 and Y848, structurally equivalent to MeV residues Y667 and L811 respectively, directly overlap with the position of the inhibitor (Supplementary Fig. 16B). The side chain of F704 adopts an orientation that intersects the compound rather than aligning parallel to its pyrazole ring, as seen for Y667 in MeV (Supplementary Fig. 16B). Additionally, H853 in RSV, corresponding to MeV H816, occupies a position that restricts F704 from flipping to a non-clashing orientation (Supplementary Fig. 16B). Another critical difference lies in residue F708 (equivalent to MeV W671), which is unable to form the hydrogen bond with the acylamino linker of ERDRP-0519 observed in MeV (Supplementary Fig. 16B). A similar steric incompatibility is observed in HMPV (Supplementary Fig. 16C). Furthermore, superposition of the LPE structure with the pre-initiation structure of VSV revealed additional clashes. Residues Y606, W609, and N692—equivalent to MeV residues L664, Y667, and T751, respectively—intrude into the binding site of ERDRP-0519 (Supplementary Fig. 16D). The catalytic residues G713, D714, and Q716 in VSV also lie within Van der Waals proximity to the inhibitor (Supplementary Fig. 16D), suggesting further steric hindrance. These clashes were similarly observed in the RABV polymerase structure (Supplementary Fig. 16E). It is worth noting that even in the predicted structure of MeV L, key residues in the fingers–palm subdomain also clash with the inhibitor (Supplementary Fig. 16F). This inconsistency highlights the limitations of structure prediction methods in accurately modeling the subtle conformational adjustments required for small-molecule binding. Consequently, prior docking studies based on such predicted models may have failed to capture the true binding mode of ERDRP-0519, leading to misleading interpretations^55^. Collectively, these structural observations, along with the lower sequence conservation in the fingers–palm subdomain among MeV, pneumoviruses (RSV, HMPV), and rhabdoviruses (VSV, RABV) (Supplementary Fig. 16G), provide a mechanistic explanation for the limited inhibitory activity of ERDRP-0519 outside the paramyxovirus family.

Although the complex structure of the MeV L–P bound to GHP-88309 could not be resolved by cryo-EM, molecular docking was performed to explore its potential binding site within the MeV L protein. Interestingly, the docking results identified a binding pocket distinct from the previously proposed site in the MeV/HPIV3 L proteins^57, 66^. Earlier docking studies suggested that GHP-88309 binds within a shallow cavity at the interface between the RdRp and PRNTase domains, but the predicted interactions were highly limited^66^. In the present study, using the high-resolution cryo-EM structure of the MeV L–P complex, we tried to re-evaluate the binding site of GHP-88309 by docking it into the ERDRP-0519 binding pocket (Supplementary Fig. 16H). The results revealed a substantially improved binding affinity at this newly predicted site (GlideScore: ∼8.45) compared to the previously reported site (GlideScore: ∼5.34)^66^, suggesting that GHP-88309 is more likely to bind at the new pocket. Notably, this alternative site partially overlaps with the ERDRP-0519 pocket and engages several of the same key residues (Supplementary Fig. 16I). The docking model suggests that the fluorobenzamide moiety of GHP-88309 is stabilized by a perpendicular π–π interaction formed between W671 and W747, In addition, Y667 and H816 engage in π–π stacking with the pyridine and benzene rings of the isoquinoline group. H816 and T751 flank the benzene ring of the fluorobenzamide, creating a clamping effect. Hydrogen bonding also contribute significantly: Q775 forms hydrogen bonds with both the nitrogen atom in the isoquinoline ring and the formamide group in the fluorobenzamide, while T748 and T751 provide additional hydrogen bonds with the same formamide group. Several other residues including T662, L664, C668, Q744, L811, and I814, contribute hydrophobic or Van der Waals interactions, further reinforcing the binding affinity and positioning of the compound within the pocket. Given its spatial overlap with the ERDRP-0519 binding pocket and its engagement with many of the same conserved residues, GHP-88309 is likely to exert its antiviral activity via a similar allosteric mechanism, i.e., likely to destabilize the active site architecture of the polymerase and thereby inhibiting RNA synthesis.

## DISCUSSION

nsNSVs rely on both the L and P proteins to assemble functional RdRP complexes for mRNA transcription and genome replication, with viral nonstructural proteins and host factors often contributing to the regulation of these two distinct processes^2, 4^. In this study, we determined two atomic-resolution cryo-EM structures of human measles virus RdRP complexes, each representing distinct functional states in the viral life cycle. In addition, we solved the cryo-EM structures of MeV and NiV RdRP bound to the non-nucleotide allosteric inhibitor ERDRP-0519 at near-atomic resolution, revealing the molecular mechanism by which this leading antiviral candidate inhibits RNA synthesis. Our structural and molecular dynamics analyses identified a conserved, druggable pocket within the RdRp domain of the L protein and demonstrated that ERDRP-0519 effectively targets both MeV and NiV polymerases. These findings underscore its potential to inhibit a broad range of paramyxoviruses and provide a blueprint for the rational design of broad-spectrum antiviral therapeutics against nsNSVs.

The mechanisms by which nsNSV polymerases coordinate mRNA transcription and genome replication remain incompletely understood and are under debate. Based on studies across multiple virus families, including paramyxoviruses, filoviruses, RSV, and VSV, two primary models have been proposed^81^. The first model derived from VSV posits that transcription and replication are carried out by distinct polymerase complexes: a “transcriptase” and a “replicase.” Both complexes are based on the L–P assembly, but the transcriptase additionally depends on several host factors^7^. Similar regulatory roles have been observed in filoviruses and pneumoviruses, where VP30 (in EBOV and MARV) and M2-1 (in RSV and HMPV) are required for transcription initiation and elongation, respectively^11, 13^. While these studies suggest that viral or host co-factors are essential for transcription, it remains unclear whether additional factors specifically regulate replication. The second model, supported by work in RSV, EBOV, and paramyxoviruses, proposes a unified polymerase that switches between transcription and replication. Transcription is initiated through a “stop-start” mechanism at the 3′ end of the genome and proceeds through sequential gene expression without encapsidation of the nascent RNA. In contrast, replication requires abundant N protein, which packages the nascent RNA co-transcriptionally, thereby protecting cis-acting signals and preventing transcription termination or mRNA modifications^2, 4, 81^.

In paramyxoviruses such as MeV, NiV and SeV, the L, P, and N alone are sufficient to drive reporter gene expression in minigenome systems, indicating that the L–P complex functions as the transcriptase. However, in RSV minigenome assays, elevated levels of N protein enhance replication but do not shift the transcription–replication balance, suggesting that N availability is necessary but insufficient to drive this switch^82^. Notably, in SeV and MeV, the nonstructural C protein has been implicated in enhancing polymerase processivity during replication and suppressing the generation of immune activating cbDI-RNAs^25, 26, 28, 29, 30, 31^. Our cryo-EM structures of MeV L–P (transcriptase) and LPC (replicase) complexes reconcile these models by revealing that C protein binding induces a replication-competent conformation of the polymerase. These structures reveal a previously unrecognized function of the C protein: in addition to antagonizing host innate immunity, C promotes a replication-competent conformation of the polymerase and facilitates N protein recruitment to the RNA exit channel. Collectively, these findings establish the C protein as a multifunctional molecular switch that regulates the transcription–replication transition, enhances polymerase processivity, and safeguards RNA synthesis fidelity. Our study unifies existing models and provides a structural framework for understanding how paramyxoviral polymerases are regulated by viral cofactors.

In paramyxoviruses, C genes are encoded only by members of the morbillivirus (e.g., MeV, CDV, RPV), henipavirus (e.g., NiV, HeV), and respirovirus (e.g., SeV, HPIV1, HPIV3) genera, but are notably absent in rubulavirus (e.g., PIV5, MuV), orthorubulavirus (e.g., PIV2), and avulavirus (e.g., NDV). This phylogenetic distribution raises an intriguing biological question: How do C-lacking viruses coordinate the transcription– replication switch and execute efficient RNA synthesis without this regulatory protein? Upon analyzing the domain architecture of P proteins across these genera, we observed a notable pattern: P proteins from C-encoding viruses tend to be substantially longer, primarily due to the presence of an extended intrinsically disordered region (IDR) between the NP-binding peptide (NPBP) and the oligomerization domain (Supplementary Fig. 17A). In contrast, C-lacking viruses encode shorter P proteins with minimal disordered regions (Supplementary Fig. 17B). This structural difference likely reflects functional adaptation. In viruses with long, disordered P proteins, the spatial constraints and low diffusion efficiency of the flexible N^0^P complex may hinder its delivery from the P protein tetramer to the nascent RNA chain. Conversely, shorter linkers may alleviate this limitation by promoting more efficient handoff of N. Our biochemical assays support this model. We found that the C protein preferentially binds monomeric or ssRNA-bound N protein rather than the N^0^P complex and is likely unable to efficiently extract the monomeric N from the P chaperone and deliver it to the elongating polymerase, thereby compensating for the functional limitations imposed by the long P-protein linker. This suggests a division of labor in C-encoding viruses, the P protein orchestrates RdRP complex assembly and RNP template engagement, while C supports N protein recruitment and enhances polymerase processivity. In C-lacking viruses, however, the P protein may perform all these roles, acting as a multifunctional coordinator of RNA synthesis. Together, these findings propose a mechanistic basis for how viruses with or without the C gene may adapt their replication strategies through structural divergence of their P proteins, highlighting the evolutionary flexibility of the paramyxoviral polymerase machinery.

To date, clinically approved antivirals that directly target essential viral enzymes remain limited, especially for highly pathogenic and lethal RNA viruses. The development of potent small-molecule inhibitors against such pathogens is therefore both urgent and technically challenging. Although prior studies have identified candidate inhibitors for morbilliviruses, including ERDRP-0519 and GHP-88309, the pharmacological basis of their binding specificity and inhibitory mechanisms has not been fully elucidated^55, 57^. In our study, we found that GHP-88309 failed to stably engage the MeV polymerase, and its predicted binding sites on MeV and HPIV3 polymerases were inconsistent^66^, suggesting a lack of conserved, specific interaction across paramyxoviruses. By contrast, our cryo-EM structure of the MeV L–P complex bound to ERDRP-0519 reveals a previously unrecognized allosteric pocket at the interface of the RdRp palm and fingers subdomains. The inhibitor inserts deeply into this cleft like a nail, with its head positioned near the catalytic center of the polymerase. Binding induces substantial conformational rearrangements in the active site, distorting its geometry and locking the enzyme in an inactive state. Importantly, this binding pocket is highly conserved across the *Paramyxoviridae* family. Despite sequence variations at the inhibitor binding site in NiV, ERDRP-0519 still binds the same pocket and induces similar inhibitory conformational changes. Furthermore, we show that the N-ethylmorpholinyl group on the piperidine ring of ERDRP-0519, originally designed to enhance solubility and oral bioavailability, extends into the central cavity of the polymerase and introduces steric clashes with the template, product RNA, and incoming NTP in the elongation state, suggesting an additional mechanism of inhibition and providing valuable insights for further optimization. Molecular dynamics simulations further support the broad-spectrum potential of ERDRP-0519 within the *Paramyxoviridae*, revealing favorable and stable binding across multiple genera. However, its binding to polymerases of other nsNSVs, such as rhabdoviruses, filoviruses and pneumoviruses, appears suboptimal, consistent with previous finding that ERDRP-0519 fails to block the replication of RSV, a distant member of paramyxovirus family^53^. Nonetheless, our structural findings define a conserved and druggable vulnerability in viral RdRPs, offering a valuable blueprint for the future design of next-generation antivirals.

Together, this study provides mechanistic insights into viral RNA synthesis regulation and presents a structural and pharmacological foundation for the development of broad-spectrum inhibitors targeting MeV, NiV, and potentially a wider range of nsNSVs. Moreover, ERDRP-0519 has demonstrated potent protective efficacy against virus-induced lethality in animal models without apparent toxicity^53, 54^, further supporting its potential for clinical development and veterinary applications.

## METHODS AND MATERIALS

### Molecular cloning

To enable efficient expression of polymerase complexes in a eukaryotic system, codon-optimized genes encoding MeV L (UniProtKB/Swiss-Prot: P35975), P (UniProtKB/Swiss-Prot: P35974), and C (UniProtKB/Swiss-Prot: P35977), as well as NiV L (UniProtKB/Swiss-Prot: Q997F0) and NiV P (UniProtKB/Swiss-Prot: Q9IK91), were individually subcloned into engineered pFastBac1 vectors (Thermo Fisher Scientific). For MeV, the full-length L protein was cloned with an N-terminal His_6_-MBP tag and a C-terminal 2×Strep-Flag tag. The full-length or truncated P protein (residues 304–507) was tagged with His_6_-MBP at N-terminus. Full-length and truncated C protein (residues 51–186) constructs were engineered with an N-terminal His_6_-MBP tag and a C-terminal GFP-Flag tag. For NiV, the truncated L protein (residues 1–1465) were subcloned with N-terminal His_6_-2×Strep-MBP tag and a C-terminal Flag tag, while the truncated P protein (residues 470–709) carried an N-terminal His_6_-MBP tag.

To measure the binding affinity between various forms of MeV monomeric N proteins and the C protein, a series of N and P protein expression constructs were generated. The apo N lacking the N-arm (residues 30–525, referred to as N_ΔN-arm_) was subcloned to a pET26b vector (Novagen) with a C terminal 6×His tag. The NPBP region of P protein (residues 1–50, referred to as P_NPBP_) was cloned into a modified pET22a vector (Novagen) with an N-terminal His_6_-MBP tag and a C-terminal SUMO tag. All constructs described above were verified by Sanger sequencing to ensure sequence fidelity.

### Protein expression and purification

To obtain well-behaved polymerase complexes suitable for structural biology studies, various combinations of MeV polymerase subunits, including full-length L with full-length P, full-length L with truncated P, and full-length L with truncated P and full-length C, were co-expressed using the Bac-to-Bac baculovirus expression system (Thermo Fisher Scientific). Similarly, NiV polymerase complex was produced by co-expression of truncated L and truncated P proteins using the same expression system.

All polymerase complexes were purified following a nearly identical procedure. Briefly, for purification of the apo L–P complexes of MeV and NiV, Sf9 insect cells (Thermo Fisher Scientific, Cat# 11496015) were harvested by centrifugation at 3,220 × g for 20 min at 4 °C, 72 hours post-infection. Cell pellets were resuspended in lysis buffer A (25 mM HEPES, pH 7.5, 500 mM NaCl, 10% glycerol, 1 mM Tris (2-carboxyethyl) phosphine (TCEP)) supplemented with an EDTA-free protease inhibitor cocktail (Selleck). Cells were lysed by sonication, and cellular debris was removed by centrifugation at 16,300 × g for 45 min at 4 °C. The clarified lysate was incubated with pre-equilibrated streptavidin affinity resin (Smart-Lifesciences) for 1 hour at 4 °C. The resin was then collected by centrifugation at 500 × g for 10 min, loaded onto a gravity flow column, and washed with 100 column volumes of buffer A. Bound proteins were eluted using buffer A supplemented with 2.5 mM desthiobiotin. Eluted proteins were concentrated using a 100 kDa molecular weight cutoff centrifugal concentrator (Millipore) and further purified by size-exclusion chromatography (SEC) on a Superose 6 Increase 10/300 GL column (Cytiva) pre-equilibrated by buffer B (25 mM HEPES, pH 7.5, 500 mM NaCl, 1 mM TCEP, and 6 mM MgCl_2_).

The MeV LPC complex was purified using the same affinity purification strategy as described above, with streptavidin resin enrichment and elution by buffer A supplemented with 2.5 mM desthiobiotin. Following concentration, the complexes were further homogenized by SEC on a Superose 6 Increase 10/300 GL column (Cytiva) in buffer C (25 mM HEPES, pH 7.5, 300 mM NaCl, 1 mM TCEP, and 6 mM MgCl_2_). To evaluate the impact of near-physiological condition (relative low salt) on protein conformation, the affinity-purified MeV L–P complex was subjected to SEC using buffer D (25 mM HEPES, pH 7.5, 150 mM NaCl, 1 mM TCEP, and 6 mM MgCl_2_).

The C protein with a C-terminal GFP tag, was expressed in Sf9 insect cells and affinity-purified using amylose resin (New England Biolabs) in buffer A. Bound proteins were eluted by buffer A supplemented with 20 mM maltose. Following ultrafiltration and concentration, the GFP-tagged C protein was further purified using a HiTrap Heparin HP column (Cytiva), eluted with a NaCl gradient between buffer E (25 mM HEPES, pH 7.5, 100 mM NaCl and 1 mM TCEP) and buffer F (25 mM HEPES, pH 7.5, 1000 mM NaCl and 1 mM TCEP). The resulting protein was concentrated and subject to SEC on a Superdex 200 Increase 10/300 GL column (Cytiva) equilibrated in buffer E.

The N_βN-arm_ and the P_NPBP_ proteins were expressed in *Escherichia coli* BL21 (DE3) cells. After transformation of the recombinant plasmids, single colonies were inoculated into 5 mL LB medium containing appropriate antibiotics and grown overnight at 37 °C. The overnight cultures were then transferred into 1 L of LB medium and grown at 37 °C until the optical density at 600 nm (OD_600_) reached ∼0.8. Protein expression was induced by addition of 0.4 mM isopropyl β-D-1-thiogalactopyranoside (IPTG), followed by incubation at 20 °C for 18 hours. Cells were harvested by centrifugation at 6,000 × g for 20 minutes at 4 °C and stored at –80 °C until purification.

To purify the N_ΔN-arm_ and the P_NPBP_ proteins, cell pellets were thawed on ice and resuspended in buffer F supplemented with an EDTA-free protease inhibitor cocktail (Selleck). Cells were lysed by sonication, and cellular debris was removed by centrifugation at 16,300 × g for 45 min at 4 °C. The clarified lysate was incubated with pre-equilibrated Ni-NTA agarose resin (Qiagen) for 1 hour at 4 °C. The resin was collected by centrifugation at 500 × g for 10 min, transferred to a gravity flow column, and sequentially washed with 100 and 10 column volumes of buffer F containing 20 mM and 50 mM imidazole, respectively. Bound proteins were eluted with buffer D supplemented with 300 mM imidazole. The eluates were dialyzed and concentrated in buffer E, then further purified using a HiTrap Capto Q Column (Cytiva), eluted with a NaCl gradient between buffer E and buffer F. Finally, proteins were concentrated and subject to SEC on a Superdex 200 Increase 10/300 GL column (Cytiva) equilibrated in buffer E. To assemble the monomeric N^0^P complex, purified N_ΔN-arm_ and P_NPBP_ proteins were mixed at a 1:1 molar and incubated on ice for 30 min prior to SEC using a Superdex 200 Increase 10/300 GL column (Cytiva) equilibrated in buffer E.

Peak fractions containing the target proteins were pooled and concentrated to 5–10 mg/mL prior to biochemical assays or cryo-EM sample preparation. Protein concentrations were determined by the absorbance at 280 nm. The concentrated samples were flash-frozen in liquid nitrogen and stored at –80 °C until use.

### Cryo-EM sample preparation

Cryo-EM sample preparation was carried out using a Vitrobot Mark IV (Thermo Fischer Scientific). All samples were diluted to a final concentration of approximately 1.2 mg/mL prior to the grid preparation. Specifically, 3 μL of MeV apo L–P complex (in 300 mM NaCl), MeV L–P complex (in 150 mM NaCl), and MeV LPC complex (in 150 mM NaCl) were applied to glow-discharged NiTi-Cu grids (1.2/1.3, 300 mesh; Guangzhou Najing Dingxin Technology). To mitigate denaturation at the air–water interface, 0.027% octyl-β-glucoside was added to the sample prior to application. For MeV L–P and NiV L–P complexes bound to ERDRP-0519, the inhibitor was add to a final concentration of 4 μM and incubated on ice for 30 min before grid preparation. Similarly, for the MeV L–P complex bound to GHP-88309, two batches were prepared with final concentrations of 60 μM and 120 μM, respectively, and incubated on ice for 30 min prior to sample application. Grids were blotted for 3.0–3.5 s at 100% humidity and 4 °C and subsequently plunge-frozen into liquid ethane. The grids were stored in liquid nitrogen until data collection.

### Cryo-EM data acquisition

Cryo-EM datasets for the MeV apo L–P complex (in 300 mM NaCl), LPC complex, and L–P complex bound to ERDRP-0519 or GHP-88309 were acquired at the Cryo-Electron Microscopy Facility of Hubei University. Micrographs were collected using a 300 kV Titan Krios transmission electron microscope equipped with a BioQuantum energy filter and a K3 Summit direct electron detector (Gatan), operated in counting mode. Automated data collection was performed using EPU software (Thermo Fisher Scientific). Datasets were acquired at a nominal magnification of 105,000 ×, yielding a calibrated pixel size of 0.851 Å. Each movie stack was dose-fractionated into 40 frames at a dose rate of 15.156 e^−^/pixel/s, with a total exposure time of 2.5 s, resulting in a total dose of ∼52.52 e^−^/Å^2^. The defocus range was set from −1.0 to −1.5 μm. A total of 14,287 (3 datasets), 14,354 (3 datasets), 4,210 (1 dataset), and 8,937 (2 datasets) movies were collected for MeV apo L–P (in 300 mM NaCl), LPC, ERDRP-0519-bound, and GHP-88309-bound complexes, respectively.

Cryo-EM data for the NiV L–P complex bound to ERDRP-0519 and MeV apo L–P complex (in 150 mM NaCl) were acquired at the Center of Cryo-Electron Microscopy, Zhejiang University, using a 300 kV Titan Krios electron microscope (Thermo Fisher Scientific) equipped with Gatan Falcon 4 detector. Datasets were collected at a magnification of 130,000 ×, corresponding to a calibrated pixel size of 0.93 Å. Each movie was dose-fractionated into 40 frames at a dose rate of 9.6 e^−^/pixel/s, with a total exposure time of 4.5 s, resulting in a total dose of ∼50 e^−^/Å^2^. The defocus range was set from −1.0 to −1.8 μm. A total of 2,510 and 5,884 (2 datasets) movies were collected for the NiV ERDRP-0519-bound and MeV apo L–P (in 150 mM NaCl) complexes, respectively. Cryo-EM data collection statistics are summarized in Supplementary Table 1.

### Cryo-EM data processing

All cryo-EM data were processed using cryoSPARC (v4.2.1, v4.5.3)^83^. Initial steps for each dataset included motion correction and contrast transfer function (CTF) estimation. For the MeV L–P complex bound to ERDRP-0519, a total of 5,846,492 particles were initially autopicked using the blob picker tool and extracted with a box size of 360 pixels. Following multiple rounds of 2D classification, 575,562 particles displaying clear structural features were selected for ab initio reconstruction. From this, a class with improved resolution was selected, and 177,478 particles were re-extracted without binning and subjected to non-uniform refinement. A final density map was obtained at 3.13 Å resolution after post-CTF refinement. Notably, although no apparent degradation of the L protein was observed, the resulting map lacked the presumed C-terminal accessory domains of L. Nevertheless, the density corresponding to the bound inhibitor in the RdRp domain was clearly visible.

For the MeV LPC complex, three datasets were independently processed for particle picking and iterative 2D classification. A total of 1,817,850 particles from the final round of 2D classification across all datasets were pooled and subjected to ab initio reconstruction into four classes. Two classes showed densities corresponding to the presumed C-terminal accessory domains of the L protein and associated C proteins. These two classes (981,463 particles) were merged for a second round of ab initio reconstruction, yielding two nearly identical classes with well-defined features. Following non-uniform and post-CTF refinement, a 2.72 Å resolution map of the LPC complex was obtained. The remaining two classes (819,802 particles), which lacked the C-terminal region of L, were merged with the apo MeV L–P dataset for further analysis (see below).

The same strategy was applied to the three datasets of the MeV apo L–P complex. After multiple rounds of 2D classification, 1,649,138 particles were selected and subjected to ab initio 3D reconstruction. One of the four classes displayed improved features and, similar to the ERDRP-0519-bound complex, lacked densities corresponding to the C-terminal accessory domains of L. This class (521,732 particles) was refined to a resolution of 2.72 Å using non-uniform refinement.

For the MeV L–P complex bound to GHP-88309, 373,200 particles were selected from two datasets and subjected to ab initio reconstruction. One class was chosen for further non-uniform refinement, resulting in a 2.88 Å resolution map. However, this map also lacked the C-terminal accessory domains of the L protein, and the density for the GHP-88309 inhibitor was not observed during model building. Given the structural similarity to the apo L–P complex map, particles from all three samples, MeV LPC complex, GHP-88309-bound, and apo, that lacking the C-terminal region were pooled and subjected to a new round of ab initio reconstruction. One of the four resulting classes, comprising 1,190,269 particles, yielded a map refined to 2.60 Å resolution.

To determine whether the absence of the C-terminal domains in L protein was due to the use of a high-salt buffer (300 mM NaCl), the apo L–P complex was reconstituted under near-physiological conditions (150 mM NaCl), and two datasets were collected. After similar processing steps, only 123,051 particles from ab initio reconstruction were selected for final refinement, yielding a map at 3.17 Å resolution. These results suggest that the apo complex in near-physiological salt conditions is less stable and more aggregation-prone, leading to a reduced yield of high-quality particles compared to high-salt conditions.

The NiV L–P complex bound to ERDRP-0519 showed greater structural stability. From only 2,510 micrographs, 468,100 particles were selected after 2D classification and subjected to ab initio reconstruction and non-uniform refinement, resulting in a 2.84 Å resolution map.

### Model building, refinement and validation

To obtain the atomic models of the MeV and NiV polymerase complexes, cryo-EM maps at atomic or near-atomic resolution were used for automated and *de novo* model building with ModelAngelo^84^. Manual adjustments were carried out in Coot^85^, followed by iterative refinement in both real and reciprocal space using Phenix^86^. Refinement was performed with artificial unit cells, electron scattering factors, and restraints on secondary structure and Ramachandran geometry. The final models were validated using MolProbity^87^.

### Microscale thermophoresis (MST) binding assay

MST was employed to quantify the binding affinity between the MeV C protein and either the L–P complex or various forms of N proteins/complexes. The C protein was C-terminally fused with eGFP and used as the fluorescently labeled target. For each interaction, 8–12 serially dilutions of the L–P complex (concentration ranging from 66.4 nM to 17 μM) or N proteins (98 nM to 200 μM) were prepared and mixed with a constant concentration (20 nM) of the labeled C protein in MST buffer (20 mM HEPES, pH 7.5, 100 mM NaCl, 1 mM TCEP and 0.05% Tween 20). After incubation in the dark at 25 °C for 30 minutes to 1 hour, 5 μL of each sample was loaded into standard-treated capillaries (Hirschmann, Germany). Measurements were performed on a Monolith NT.115 instrument (NanoTemper Technologies, Germany) using blue/red filters at 25 °C. The excitation settings were 40% blue LED power (excitation 460–480 nm, emission 515–530 nm) and 60% infrared laser power, with thermophoresis cycles set to 3 s off and 20 s on. Each binding isotherm was generated from 8–12 titration points and repeated in three independent experiments. Data analysis was conducted using NanoTemper analysis software, and dissociation constants (*K*_d_) were calculated from saturation binding curves at equilibrium.

### Molecular docking

ERDRP-0519 (PubChem CID: 57521469) and GHP-88309 (PubChem CID: 50962592) were prepared for molecular docking studies using the LigPrep tool of the Schrödinger suite v.2021-2 (Schrödinger). The compounds’ SDF files were converted into energy-minimized 3D conformations using the OPLS3e force field. Hydrogen atoms were added, and all possible ionization states and stereoisomers were generated at a pH of 7.0 ± 0.5 using Epik, with the desalt option enabled. The structures of L proteins from PIV3 (PDB: 6V85), PIV5 (PDB: 8KDB), MuV (PDB: 8IZL), NDV (PDB: 7YOU), and MeV (cryo-EM structure determined in this study) were processed using Maestro Schrödinger’s Protein Preparation Wizard, which involved adding missing hydrogen atoms, assigning bond orders, optimizing hydrogen bonding networks, and removing water molecules located more than 5 Å away from protein residues. Subsequently, the protein structures were optimized by predicting the pKa values of ionizable residues with PROPKA, followed by restrained energy minimization using the OPLS3e force field. Receptor grids (size 10 × 10 × 10 Å) were generated at the predicted catalytic cavities or central channels using Glide’s Receptor-Grid-Generation tool. Compounds were docked into these grids using Schrödinger’s Ligand Docking tool, applying a van der Waals radii scaling factor of 0.8 for non-polar atoms and a partial charge cut-off of 0.15. The initial docking was performed using the standard precision protocol, followed by a more rigorous extra precision protocol with flexible ligand sampling to refine the results.

### Systems preparation for MD simulations

We constructed six simulation systems of L/ERDRP-0519 complexes from MeV, NiV, HPIV3, PIV5, MuV, and NDV. For the MeV and NiV systems, cryo-EM structures in this study were applied. For the HPIV3, PIV5, MuV, and NDV systems, the docked complex structures of ERDRP-0519 with the corresponding L proteins were used as the initial coordinates. All six L/ERDRP-0519 complexes were solvated using the Solution Builder module in CHARMM-GUI. Each system was placed in a rectangular TIP3P water box (box dimensions: 120 × 120 × 120 Å; crystal type: CUBIC; α = β = γ = 90°) and neutralized with 0.15 M NaCl. The final systems contained about 160,000 atoms, depending on protein size and composition.

### Molecular dynamics (MD) simulations

All-atom molecular dynamics simulations were performed using the GROMACS 2024.1 package with the CHARMM36m force field^88^. Parameters for ERDRP-0519 were assigned using the CGenFF server^89^. Each system was first subjected to energy minimization using the steepest descent algorithm. Two rounds of equilibration were run under both isothermal–isochoric and isothermal–isobaric conditions, with gradually decreasing position restraints applied to heavy atoms of the proteins and ERDRP-0519. After releasing all restraints, production simulations were carried out. Simulations were performed at 303 K and 1 atm using the V-rescale thermostat and C-rescale barostat. Bond lengths were constrained using the LINCS algorithm, allowing a 2-fs integration timestep. A 12.0 Å cutoff was applied to short-range van der Waals and Coulombic interactions, and long-range electrostatics were treated using the particle mesh Ewald (PME) method. Structural dynamics and ligand-protein interactions were analyzed using GROMACS built-in tools.

### Figure preparation

Protein-protein and protein-small molecule interaction analysis were performed using PyMOL (https://pymol.org/2/) and PDBePISA (https://www.ebi.ac.uk/pdbe/pisa/). Structural figures and electron density map visualizations were prepared using UCSF Chimera^90^, ChimeraX^91^ and PyMOL. Multiple sequence alignments were generated using Multalin^92^ and ESPript^93^. The structures of the MeV L–P complex and C protein dimers from CDV, RPV, NiV, HeV, HPIV1, HPIV3, and SeV were predicted using the AlphaFold3 server^94^ (https://alphafoldserver.com/) for structural comparison with the experimentally determined models. Conservation analysis of the ERDRP-0519 binding pocket across paramyxoviral polymerases was conducted using the ConSurf server^95^ (https://consurf.tau.ac.il/consurf_index.php).

## Data availability

The authors declare that all data supporting the findings of this study are available in the article, its Supplementary Information, and its Source Data. The cryo-EM maps and atomic coordinates generated in this study have been deposited to the Electron Microscopy Data Bank (EMDB) and the Protein Data Bank (www.PDB), respectively. The accession codes are as follows: EMD-65364 and PDB: 9VUI for MeV LPC complex; EMD-65365 and PDB: 9VUJ for MeV apo L–P complex in 300 mM salt buffer; EMD-65366 and PDB: 9VUK for MeV apo L–P complex in 150 mM salt buffer, EMD-65367 and PDB: 9VUL for MeV L– P complex bound to the allosteric inhibitor ERDRP-0519; and EMD-65368 and PDB: 9VUM for NiV L–P complex bound to ERDRP-0519.

## Acknowledgements

We thank the staff at the Core Facility at the Institute of Life Sciences (LSI), Zhejiang University, for providing instrumentation and technical support, particularly Ms. Weina Shang and Ms. Jie Ma for their assistance and guidance in instrument operation. We are also grateful to the staff of the Cryo-Electron Microscopy Facility of Hubei University and the Center of Cryo-Electron Microscopy (CCEM) at Zhejiang University, with special thanks to Dr. Shenghai Chang for his support in cryo-EM data collection. This work was supported by the National Natural Science Foundation of China (32371344 to H.R. and 32471319 to S.W.), the startup funding from LSI, Zhejiang Key Laboratory of Molecular Cancer Biology at Zhejiang University, as well as the National Young-Thousands Talents Program. Additional support was provided by the Distinguished Young Scholars of Hubei Province (2022CFA078 to S.W.), and Knowledge Innovation program of Wuhan-Shugung Project (2023020201020418 to S.W.).

## Author Contributions

H.R. conceived and supervised the study. T.D., G.L., K.J., and Y.C. performed molecular cloning, protein expression, purification, and cryo-EM sample preparation. J.W., T.D., and S.W. collected the cryo-EM data. H.R. and T.D. determined the cryo-EM structures, built and refined the atomic models. X.Z. provided small-molecule inhibitors for structural studies and contributed to cryo-EM data processing. T.D. and R.X. performed MST measurement and data processing. C.Y. and Q.Z. performed molecular docking and MD simulation study. H.R., D.T., and Q.Z. analyzed the data and wrote the manuscript. G.S. and L.Z. reviewed the manuscript and contributed valuable suggestions for its revision. H.R., S.W., Q.Z., and T.D. participated in discussion and revised the manuscript.

